# Intersegmental transfers drive target search in an RNA-targeting CRISPR system

**DOI:** 10.64898/2026.06.18.732892

**Authors:** Ofer Kimchi, Benjamin B. Larsen, Emily Gibson, Aartjan J. W. te Velthuis, Cameron Myhrvold

## Abstract

Sequence-specific RNA-binding proteins (RBPs) must efficiently locate their targets among a multitude of cellular RNAs. Cas13, an RNA-guided CRISPR protein, represents an ideal model system in which to study this search process. Cas13 combats bacteriophage infection by cleaving RNA nonspecifically upon binding of its crRNA to the target RNA sequence; thus, Cas13’s search for its RNA target comes with a time constraint determined by the rate of phage multiplication. The mechanism by which Cas13 locates its target within this critical window remains unknown. Here, we investigate Cas13’s mechanism of target search through integration of biophysical modeling, activity assays, and biochemical characterization. We show that Cas13 employs facilitated diffusion to accelerate its search, and find that Cas13’s search time when targeting RNAs of different lengths cannot be explained by 1D sliding, the search mechanism used by many DNA-binding proteins. We propose that Cas13 primarily searches for its RNA target by intersegmental transfers (ITs), non-specifically binding the RNA at two locations and directly switching between them without fully dissociating from the RNA. We develop a biophysical model for ITs in an RNA context that we subsequently validate experimentally. Furthermore, we demonstrate that ITs can differentially accelerate the search process for a broad class of RNA-binding proteins, as opposed to their DNA-binding counterparts, due to RNA’s short persistence length and the heterogeneity of RNA lengths in the cell. Our results illuminate how Cas13 achieves rapid target recognition in a complex RNA environment, and implicate ITs as a potentially widespread solution to the RNA search problem.

## INTRODUCTION

Sequence-specific nucleic acid binding proteins must efficiently search vast physical and sequence space to locate their targets on a biologically relevant timescale. The search processes of RNA-binding proteins (RBPs) in particular present a unique challenge compared to their DNA-binding counterparts. First, the scale of the search is substantial: for example, an *E. coli* cell contains hundreds of thousands of unique RNA molecules (Fig. 1A) including an estimated ∼10 times more non-ribosomal RNA nucleotides than DNA base pairs^1–4^. Moreover, these RNA molecules have a large length distribution, ranging from under 50 nt for the shortest sRNAs to well over 4 kb for the longest mRNA species, in addition to wide variation in features such as GC content and secondary structure^5^.

**Figure 1:**
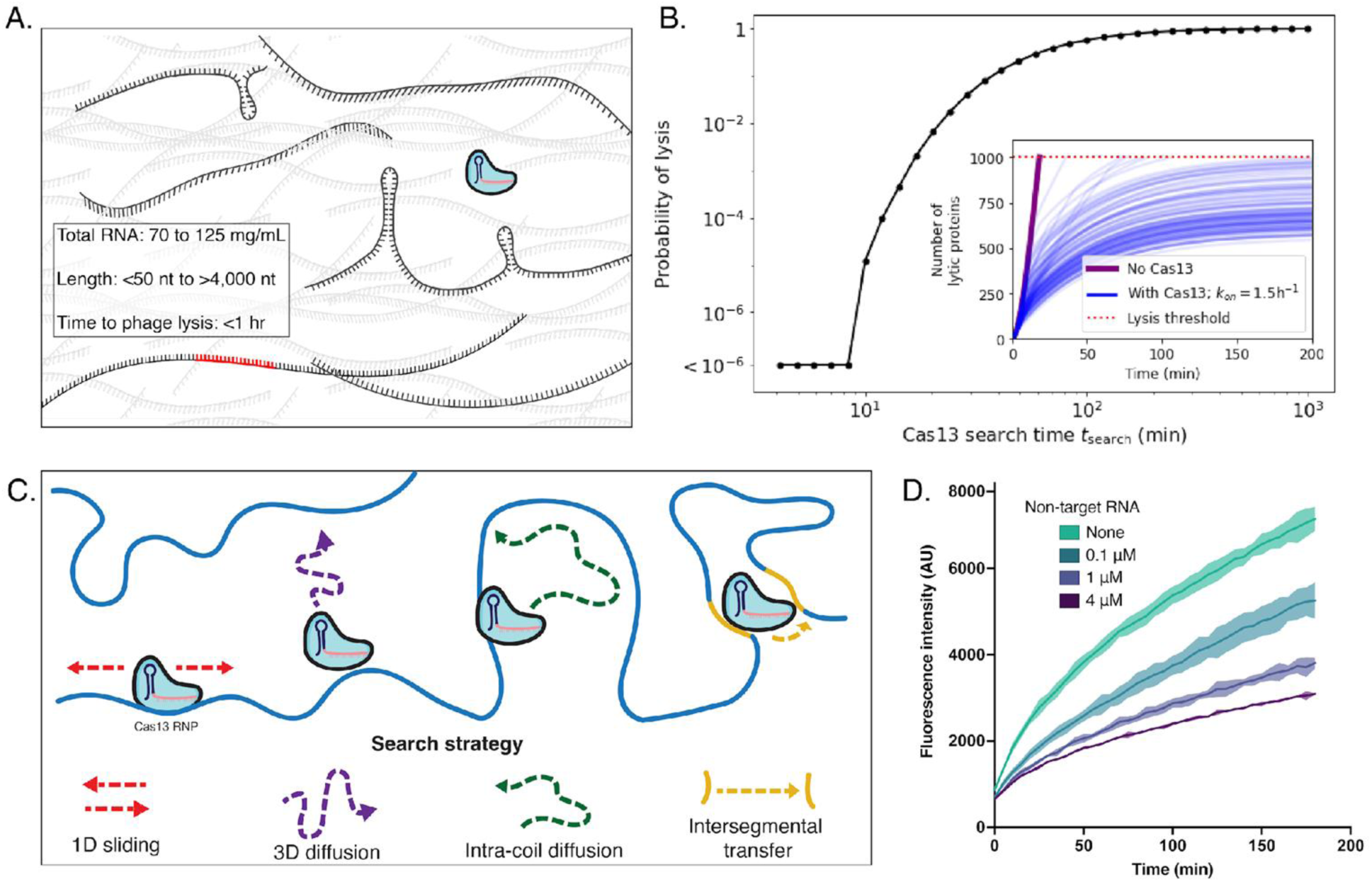
Cas13 efficiently searches an expansive RNA sequence space. **A.** Cas13 must search through abundant cellular RNAs of varying lengths and secondary structures to locate a 20-28 nt protospacer sequence on a relatively short timescale. **B.** Inset: Stochastic trajectories of the accumulation of lytic proteins for a given system with *k*_on_=(*t*_search_)^-1^=1.5/h. Main figure: Fraction of stochastic trajectories resulting in cell lysis, as a function of Cas13 search time *t*_search_. Model is described in Section S5. **C.** Visual summary of four previously described search mechanisms that a sequence-specific RNA- or DNA- binding protein may utilize to find its binding partner. **D.** Fluorescence kinetic curves show that addition of increasing amounts of single-stranded non-target RNA leads to slower Cas13 activation *in vitro*. Target RNA concentration: 0.17 nM. Error of n = 2 technical replicates is shown by shading.

CRISPR-Cas13 represents an effective model system for studying protein-RNA interactions, including the RNA search process. Cas13 is an RNA-guided, RNA-targeting CRISPR nuclease which has been widely adopted for technological applications including viral diagnostics, RNA knockdown, and RNA imaging^6–10^. Upon binding its RNA target, Cas13 nonspecifically cleaves other RNAs in solution, enabling a clear readout of Cas13 activity in different environments through the addition of short quenched fluorescent RNA reporters.

Cas13’s canonical function in bacteria is to provide immunity from bacteriophage infection by sequence-specifically binding to phage mRNAs and subsequently unleashing nonspecific RNA cleavage, which induces cell dormancy and terminates phage infection prior to cell lysis^11,12^. In the absence of phage defenses such as Cas13, a lytic infection leads to infection-competent progeny within as few as 30 minutes for many phages, at which point it is likely too late to initiate an effective immune response^13,14^. Cas13 must locate its target RNA quickly enough to mount a successful response during that window, and thereby prevent a runaway phage infection and collapse of the bacterial population (Fig. 1B).

To speed their search beyond the rate set by 3D diffusion, nucleic acid binding proteins typically employ a process termed facilitated diffusion: diffusing in 3D until binding non-specifically to a section of DNA or RNA, then searching locally on the molecule, before dissociating and repeating the process^15–17^. Several methods for local search have been hypothesized based on studies of DNA-binding proteins: 1D sliding, in which the protein binds nonspecifically to DNA and moves randomly along the coil before dissociating; intracoil diffusion (also sometimes referred to as “hopping” and “jumping”) in which the protein dissociates from the DNA and diffuses in 3D within the coil before reassociating; and intersegmental transfers (ITs), in which the protein utilizes multiple non-specific binding patches to directly transfer between different DNA segments without fully dissociating (Fig. 1C)^17–19^. The presence or absence of multiple nonspecific binding patches is the key difference between intracoil diffusion and ITs, as the former involves multiple rounds of dissociation and reassociation of the protein from the polymer, while in the latter scheme the protein remains (non-specifically) bound to the polymer throughout the search process. As a result, intracoil diffusion is almost entirely driven by the polymer itself, and is largely invariant to properties of the protein. On the other hand, ITs are driven by both polymer properties and protein properties, and thus can be evolutionarily tuned in different ways by different proteins, just as the distance of 1D sliding can be tuned evolutionarily^20,21^.

Facilitated diffusion has been observed in numerous DNA-binding proteins, including CRISPR-Cas9 and Cas12 whose local search is dominated by 1D sliding^21–26^. Though far less studied than their DNA counterparts, the search processes of certain RBPs have also been investigated. Recent work on the RBPs Hfq, RNAse E, and Argonautes have argued that they perform 1D sliding as part of their target search^27,28^. Another study on Argonautes argued that intracoil diffusion plays a dominant role^29^. To our knowledge, in none of these studies of RNA-binding proteins were intersegmental transfers explicitly considered as a potential search strategy, nor have quantitative arguments been made for the circumstances in which different facilitated diffusion strategies may be preferred for an RNA search.

Here, we explore how sequence-specific RBPs can rapidly find their targets within the massive and heterogeneous landscape of RNA molecules in the cell. We use Cas13 as a model RBP for our study since, beyond its applications in basic research and biotechnology, its highly active *trans*-cleavage activity provides a simple fluorescent readout for target binding and efficient multiplexing of experimental conditions^30,31^. Coupling Cas13 binding and activity assays together with biophysical modeling, we demonstrate that Cas13 utilizes facilitated diffusion to accelerate its search beyond the limit imposed by 3D diffusion. Furthermore, we show that Cas13 uses ITs as the dominant search modality, and argue that ITs can be more efficient than other facilitated diffusion strategies for a broad class of RNA-binding proteins, especially those targeting long RNAs (≳1,000 nucleotides in length).

## RESULTS

### Non-target RNAs slow Cas13’s search

A major search obstacle faced by RNA-binding proteins, in comparison to DNA-binding proteins, is the quantity and heterogeneity of RNA molecules in solution. We first sought to establish whether the presence of RNAs lacking the 28-nt target or “protospacer” sequence (referred to hereon as non-target RNAs) substantially affects Cas13 search kinetics. To quantify our expectation for this effect, we consider the rate of target binding, *k*_on_, for a searching protein which is initially not bound to the target RNA. This rate is equal to the inverse of the average search time *τ*_search_, and has previously been found by several groups^17,20,32,33^ (also, see Supp. Section S1) to scale with system properties as

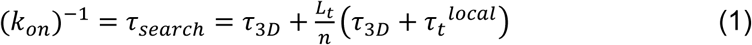

where *τ*_3D_ is the time it takes the protein to bind non-specifically to some section of the target RNA, *L*_t_ is the number of possible binding sites in the target RNA (one of which is the protospacer, i.e. the target site), *n* is the number of sites typically searched upon each non-specific binding event (and is >1 for facilitated diffusion) and *τ*_t_^local^ is the time taken to search those sites. In a solution with only target RNAs (i.e. RNAs containing the protospacer), *τ*_3D_ is given by the inverse of the Smoluchowski rate *k*_t_^3D^=4π*D_t_r_t_c_t_* , where *D*_t_ is the sum of the diffusion coefficients of the protein and target, *r*_t_ is the radius of gyration of the target RNA, and *c*_t_ is the target RNA concentration. However, in the presence of non-target RNAs, this rate is modified to be

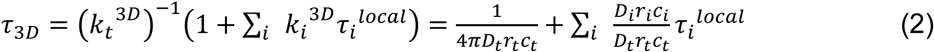

where terms with the *i* subscript denote the *i*^th^ species of non-target RNA in solution. Eq. (2) quantifies how the presence of non-target RNAs will slow down the search process.

To measure this effect experimentally in our model system, we utilized a Cas13 activity assay in which Cas13’s activation by target binding leads to the trans-cleavage of a quenched fluorescent reporter molecule. The rate of the resulting fluorescence increase is referred to as “activity” and labeled *ξ*. As fluorescence assays using *in vitro* transcribed RNA tend to exhibit minimal variation (coefficient of variation <5%), we used two technical replicates for most activity assays throughout^34–36^. We measured LwaCas13a activation kinetics in the presence of increasing concentrations of non-target RNA, while keeping the target RNA concentration constant. We designed both target and non-target RNAs to be single-stranded to control for structure effects (see Methods). We find that increasing concentrations of non-target RNA reduce Cas13 activity (Fig. 1D). Since no element of the target RNA was changed, this establishes that Cas13 activity can be used as a proxy for the search time kinetics modeled by Eq. (1); although the precise relationship between the two is not established here, we make use of the fact that *ξ*(*k*_on_) is a monotonically increasing function throughout this work. An alternative hypothesis that non-target RNA predominantly affects Cas13’s *trans*-cleavage rather than its search time is not supported by follow-up experiments (see Supplement, Section S2). Notably, Cas13 is still activated to a significant degree even when the target RNA is outnumbered by non-target RNAs by a factor of over 20,000.

### Cas13 employs facilitated diffusion in its search

We next sought to establish whether Cas13 uses facilitated diffusion in its search process. We reasoned that if Cas13 uses facilitated diffusion, the Cas13-crRNA complex may locate its target RNA faster than the crRNA alone, which must search solely through 3D diffusion. We designed a strand displacement assay to directly report target binding independent of Cas13 activity (Fig. 2A). We found that fluorescence kinetics were ∼3x faster in the presence of catalytically inactive Cas13 (dCas13) compared to the crRNA alone, despite Cas13’s larger size and slower diffusion (Fig. 2B-C)^37^.

**Figure 2:**
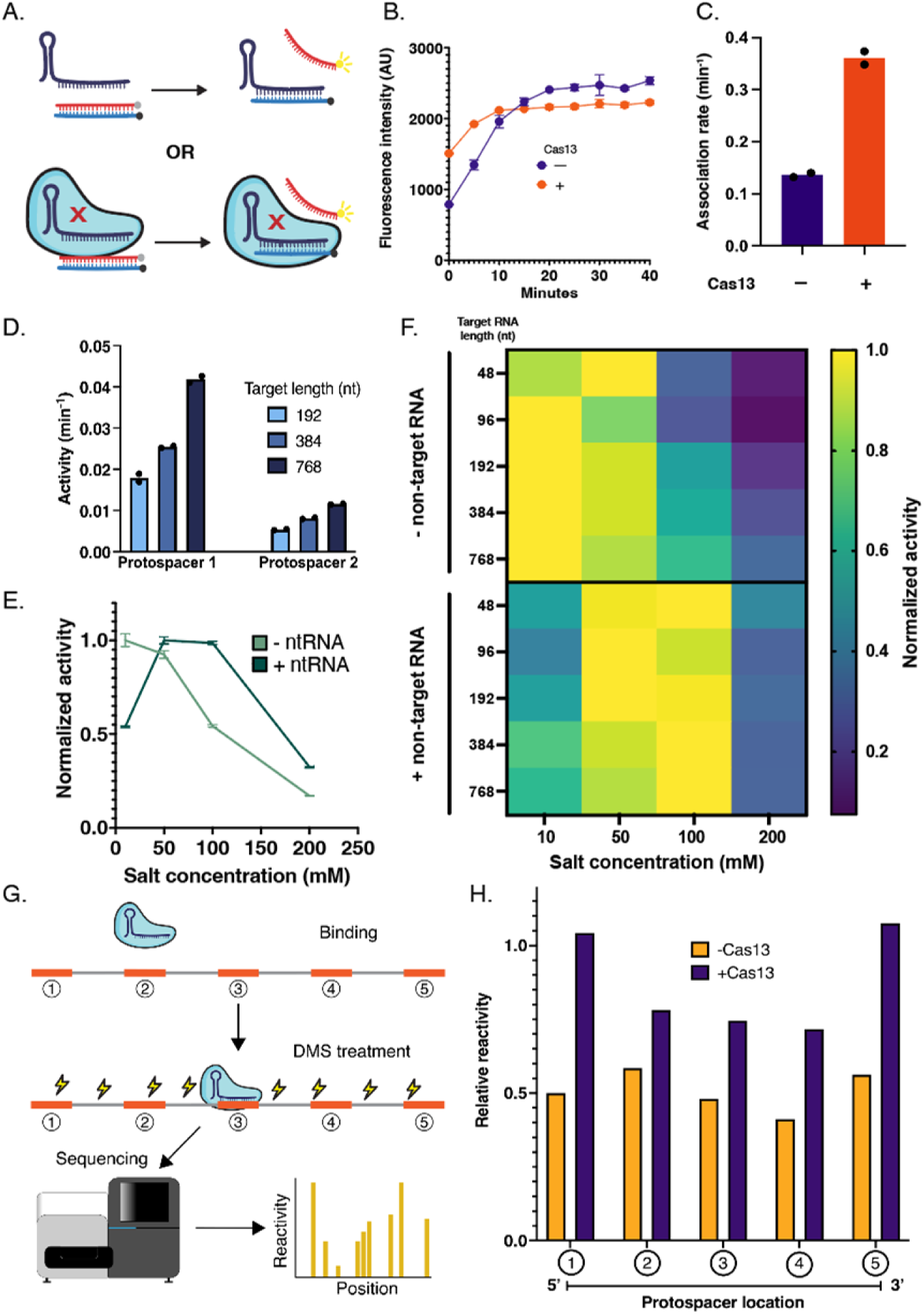
Cas13 locates its RNA target by facilitated diffusion. **A.** Schematic of an *in vitro* assay which directly reports kinetics of target RNA binding with or without the presence of catalytically inactive (dead) Cas13 via a strand-displacement readout; red X indicates dead Cas13. **B.** Fluorescence kinetic curves reveal faster target binding in the presence of dead Cas13 compared to the complementary crRNA alone. Error bars represent the standard deviation of two technical replicates. **C.** Activity quantification (pseudo-first order rate constant) of the kinetic curves shown in B. **D.** Activity quantification showing an increase in Cas13 activity when targeting RNAs of increasing length, for two unique protospacer sequences. **E.** Cas13 activity profile across different salt concentrations in an *in vitro* assay in the presence or absence of RNA for a 192 nt target RNA. Error bars represent the standard deviation of two technical replicates. **F.** Heatmap showing Cas13 activity across varying salt concentrations for target RNAs of different lengths in the presence or absence of RNA. Y axis markings indicate target RNA length. **G.** Schematic of a DMS Cas13 footprinting assay using a target RNA containing five identical protospacer sequences. In brief, the binding of the crRNA to the target RNA protects the bound region from DMS-induced mutagenesis, which can be detected with a sequencing readout and used to construct a crRNA occupancy profile. **H.** Relative DMS reactivity of the five different protospacer sequences in the presence or absence of Cas13, highlighting Cas13-mediated binding bias toward the center of the molecule.

To further probe whether Cas13 uses facilitated diffusion to speed up its search, we examined the scaling behavior of the search rate *k*_on_ as a function of target length, *L*_t_. Changing *L*_t_ affects *k*_on_ in two different ways: first, directly (Eq. 1) as longer targets have more non-target sites which Cas13 must search before finding the protospacer; second, indirectly (Eq. 2) as longer targets are physically larger, leading to faster non-specific binding to the molecule (i.e. increasing *k*_t_^3D^). For a pure 3D search, the former effect is always dominant, such that increasing the target length will slow down the search (see Supp. Section S2). However, that is not necessarily the case for a search using facilitated diffusion. For small enough target RNAs, a protein employing facilitated diffusion may display an increase in *k*_on_ as *L*_t_ is increased^18^. We find that Cas13 activity (and therefore also *k*_on_) increases with *L*_t_ for target RNAs between 192 and 768 nt. This scaling behavior was consistent across two different protospacer sequences, despite variations in absolute activity between them (Fig. 2D, Supp. Fig. 1).

Another hallmark of facilitated diffusion concerns the effect of ionic conditions on the search time. Lower salt concentrations are expected to increase the dwell time of a (positively-charged) protein on a (negatively-charged) polymer such as RNA. Therefore, in the absence of facilitated diffusion, i.e. for *n*=1, decreasing the salt concentration will simply increase *τ*_t_^local^, and thereby decrease *k*_on_. However, a search employing facilitated diffusion can exhibit a regime in which decreasing the salt concentration will increase *k*_on_ due to an increase in *n*. We find that Cas13 activity monotonically increases with decreased [NaCl] under the conditions tested (Fig. 2E; light green). To control for potential effects of salt on the equilibrium Cas13 activity, we tested the effects of changing [NaCl] in the presence of non-target RNA. With non-target RNA present, low values of [NaCl] are expected to lead Cas13 to dwell longer on non-target RNAs, increasing *τ*_i_^local^, and thereby decreasing *k*_on_. In agreement with this prediction, we find a non-monotonic dependence of Cas13 activity on [NaCl] in the presence of non-target RNAs (Fig. 2E; dark green). Across different target RNA lengths, the maximum Cas13 activity was consistently found at a higher salt concentration in the presence of non-target RNAs than in their absence (Fig. 2F, Supp. Fig 1).

Finally, we explored the effects of different protospacer placements within the target molecule. While a search using pure 3D diffusion will tend to find any protospacer location with equal probability, a process employing local search with correlated motion between sites searched will preferably find protospacers located nearer the center of the RNA^38^. To observe potential preferences in Cas13’s target binding across the length of an RNA molecule, we created a novel methylation-based footprinting assay based on dimethyl sulfate mutational profiling with sequencing (DMS-MaPSeq) using dCas13^39,40^. We designed a 254 nt-long unstructured target RNA containing five identical protospacer sequences equally spaced throughout the molecule (Methods). The dCas13-crRNA complex or crRNA alone was then allowed to bind to the target RNA. The RNA was then treated with DMS, a strong methylating agent which acts on unpaired adenine and guanine bases, resulting in a methylation profile corresponding to the unbound regions of RNA and leaving behind unmethylated “footprints” where dCas13-crRNA or crRNA alone was bound (Fig. 2G). A highly processive reverse transcriptase converts these methylation marks to mutations in the resulting cDNA, which can then be analyzed by next-generation sequencing to establish a binding profile. We found that while the crRNA alone has a uniform binding profile across the five protospacer sequences, the dCas13-crRNA complex is disfavored from binding to the two protospacers at the 5’ and 3’ termini of the RNA as evidenced by the elevated rate of DMS reactivity within these protospacers (Fig. 2H, Supp. Fig. 2). Taken together, the dependence of Cas13 activity on target length, effects of ionic concentration, and observed edge effects provide strong evidence of facilitated diffusion by Cas13 in its target search process.

### Scaling of search time with target length can distinguish between search strategies

We next asked which facilitated diffusion strategy Cas13 uses in its search. As the search process for several other RBPs has been found to involve 1D sliding^27,41^, and given the prevalence of the 1D sliding strategy in DNA-binding proteins including other CRISPR proteins^22–26^, we initially hypothesized that Cas13 would employ 1D sliding in its search process.

A sliding based search has the property that *τ*_t_^local^=*n*^2^/2*D*_1D_ (with *D*a 1-dimensional diffusion constant) is independent of *L*_t_, while *τ*_3D_∼(*L*_t_)^-*ν*^, where *ν*<1 is the Flory exponent (*ν*=½ for a random walk) (see Section S2 for a more detailed calculation). The overall target acquisition rate for a sliding-based search is then given by

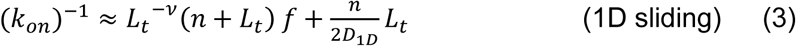

where *f* is independent of *L*_t_. Therefore, in the absence of non-target RNAs, the overall relationship of *ξ*(*L*_t_) is expected to be a non-monotonic concave function, as has been noted in other work^18^.

An alternative search strategy would be intracoil diffusion. Intracoil diffusion is a qualitatively different form of facilitated diffusion because it is unclear to what extent the protein itself can be evolutionarily tuned to modulate its behavior. Prior work has shown that intracoil diffusion plays a dominant role in Argonautes’ search (though we note the authors of this paper used the phrase “intersegmental transfer” to refer to this process; this should not be confused with intersegmental transfers as they are defined in our work and elsewhere)^29^. Intracoil diffusion is typically separated into two qualitative regimes: short “hops” of distance <*b* wherein the protein binds to a spot correlated with its previous binding position, and longer “jumps” of distance ≫*b* with no such correlation^32,42^. Given the short persistence length of RNA, we neglect hops in this work. As derived by Halford and Marko with simple scaling arguments, for intracoil diffusion, *n*∼(*L*_t_)^1-*ν*^ ^20^. Assuming that the time spent diffusing intramolecularly is negligible, and assuming a uniform dwell time *τ*_d_ for each non-specific binding event, a search dominated by intracoil diffusion has:

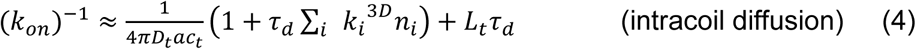

where *a*=0.33 nm is the length of each site of the RNA (see derivation in Supp. Section S2). In other words, intracoil diffusion enables the search process to largely behave as if the target site is on its own, separate, RNA strand. In the absence of non-target RNA, Eq. 4 implies that a search dominated by intracoil diffusion would exhibit an association rate that decreases monotonically with *L*_t_.

### Neither 1D sliding nor intracoil diffusion are the dominant search strategy used by Cas13

Eqs. 3 and 4 demonstrate that studying the qualitative properties of *ξ*(*L*_t_) could enable us to distinguish between different search strategies. For example, finding that Cas13 activity scales with target length as a non-monotonic concave function would rule out intracoil diffusion as the dominant search mechanism. We therefore designed three sets of unstructured target RNAs with lengths ranging from 48 to 768 nt, containing the same protospacer sequence but different flanking sequence contexts. For all target sequences, measured Cas13 activity in the absence of non-target RNA was non-monotonic and convex—initially high at very short target lengths before a decrease and subsequent increase in activity as *L*_t_ was increased (Fig. 3A, Supp. Fig 3). This qualitative behavior cannot be explained by either Eq. 3 (1D sliding; green curve in Fig. 3B) or Eq. 4 (intracoil diffusion; yellow curve in Fig. 3B).

**Figure 3:**
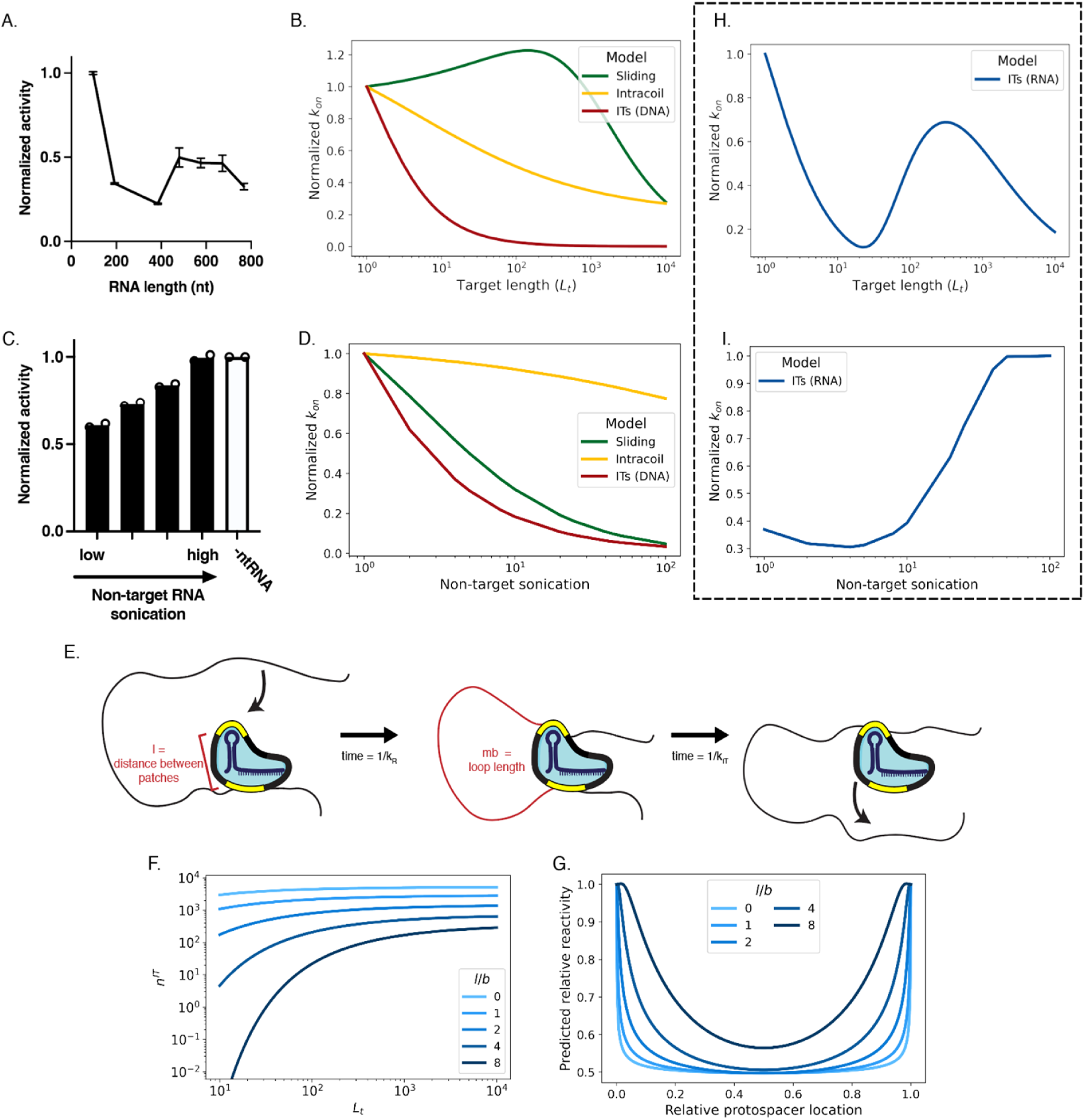
Observed effects of RNA length on Cas13 activation are consistent with intersegmental transfer, but not with other models. **A.** Cas13 activity profile when targeting RNAs of varying lengths containing a protospacer in the middle of the molecule. Error bars represent the standard error of two separate experiments with two technical replicates each. **B.** Modeling predictions of the rate of target binding as a function of RNA target length for different search strategies. **C.** Cas13 activity as a function of average non-target RNA length, generated by spiking target RNA into solutions containing a constant concentration of non-target RNAs that were subjected to different amounts of sonication. **D.** Modeling predictions of the rate of target binding as a function of the degree of non-target RNA sonication for different search strategies. **E.** Schematic illustrating the intersegmental transfer process for Cas13. **F.** The predicted number of ITs performed is plotted as a function of RNA length; colors correspond to the distance between binding . **G.** An IT-based search model predicts edge effects in binding, where sites near the middle of the RNA are searched more than sites near the edges. **H.** The predicted effect of target length on the rate of target binding for an IT-based search. I. The predicted rate of target binding as a function of the degree of non-target sonication for an IT-based search.

To further explore the role of RNA length on Cas13 activity, we took ∼1000 nt-long non-target RNAs subjected to different amounts of sonication. We then spiked in a constant concentration of target RNA, thereby creating samples with constant values of *L*_t_ and constant amounts of non-target RNA nucleotides, but with differing length distributions for the non-target RNA (Fig. 3C, Supp. Fig. 3). The most highly sonicated sample contained a large number of shorter non-target RNA molecules and the less-sonicated samples contained fewer and longer non-target RNA molecules. Following a similar analysis to Eq. 3 (see Supp. Section S2), a search dominated by 1D sliding would exhibit faster association for lower amounts of non-target RNA sonication, as only the number of non-target RNAs, and not their lengths, affect the search time (green curve in Fig. 3E). Similarly, Eq. 4 indicates that the behavior of a search dominated by intracoil diffusion is expected to be largely invariant to sonication for small amounts of sonication, and exhibit similar qualitative behavior as that predicted for sliding for large amounts of sonication (yellow curve in Fig. 3D). However, we once again see behavior inconsistent with both of these predictions: higher Cas13 activity for *higher* amounts of non-target RNA sonication (Fig. 3C).

### A model for intersegmental transfers (ITs)

Having found that Cas13’s observed behavior is incompatible with both a search dominated by 1D sliding and one dominated by intracoil diffusion, we consider a third possibility, that of intersegmental transfers (ITs). ITs can be performed when the protein has more than one patch capable of associating non-specifically with the RNA. A segment of the RNA non-specifically associates to the protein at one of these patches, and while that segment remains bound, stochastic thermal motion brings a second region of RNA to bind transiently to the second patch. If the first RNA segment dissociates before the second, the protein has effectively transferred between regions of RNA which can be quite distant along the RNA contour, but which were transiently near each other in 3D space (Fig. 3E). As an analogy, consider the protein as a passenger at a railway transit station; it can switch between different train lines (different RNA segments) without leaving the station.

The generic nature of these non-specific RNA-binding patches on the protein is a key point. In prior work, ITs have typically been considered for multimeric proteins, which have multiple sites able to bind *specifically* to the target^43^. However, this need not be the case. In order to perform ITs, an RBP need only be capable of associating non-specifically with two RNA segments simultaneously. We refer to the locations for non-specific association as “binding patches” on the protein, but these may be as simple as a patch of positively-charged residues on the protein surface, capable of non-specifically attracting negatively-charged RNA.

Existing models for ITs on the DNA cannot explain the behaviors observed in the experiments shown in Fig. 3 (red curves in Fig. 3B and 3D); however, these existing models are not necessarily appropriate for a search on the RNA. For example, prior work has indicated that ITs can particularly speed up the search on the DNA when the protein rarely or never dissociates from the polymer^44,45^; such a search would be highly inefficient in the context of RNA given the prevalence of non-target RNAs. Importantly, though, prior models have not accounted for the relationship between the rate of performing ITs and the overall dissociation rate of the protein from the polymer, nor have they considered the effects of a short polymer persistence length compared to protein size, as is the case for RNA^17,44–47^. Therefore, motivated by our findings that Cas13 uses a facilitated diffusion strategy that is neither sliding nor intracoil diffusion, we decided to revisit this problem and develop a model for an IT-based RNA search.

### Short ITs on RNA are exponentially suppressed

The first question we ask is how quickly each IT can be performed. ITs are mediated by the dynamic fluctuations of the RNA, which have previously been found to be well described by the Zimm model^48^. If a section of RNA is bound to one of the non-specific binding patches, the rate at which a region of RNA a contour length of *m* Kuhn lengths away binds to the other binding patch, located a distance *l* away in 3D space, scales approximately as

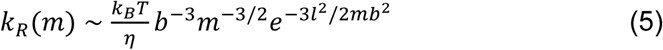

where *b* is the Kuhn length of the RNA (∼2 nm^49^; equal to twice the persistence length), *k*_B_ is Boltzmann’s constant, *T* is temperature in Kelvin, *η* is the viscosity of solution, and for clarity we have omitted a numerical prefactor as well as a term logarithmic in the reaction radius^50,51^. The heavy-tailed scaling of *k*_R_∼*m*^-3/2^ in Eq. 5 is a key feature of ITs: they are superdiffusive, and enable the protein to search far from the site of initial binding, in contrast to diffusive 1D sliding which is better suited to short-distance searches. Specifically, ITs are an example of a Lévy flight, and the mean step size of an IT scales with the total length of the RNA. Because the Kuhn length of DNA is ∼100 nm^52^ and protein diameters are typically much smaller, *l* ≪ *b* when considering a DNA search and the exponential term has been neglected in prior models of ITs for DNA-binding proteins. However, this exponential is non-negligible for RNA-binding proteins, and can have a substantial impact on the search, as we will show.

*k*_R_ gives the rate at which RNA strands encounter the protein, but the rate at which the protein actually performs an IT is given by the dissociation rate of each binding patch: ITs are performed when one binding patch dissociates and rebinds before the other dissociates. We therefore argue that the rate of performing ITs and the overall dissociation rate of the protein from the polymer must be coupled; this coupling is the second major deviation of our work from prior models of ITs. Here, we assume for simplicity that the dissociation rates of the two binding patches are equal and given by *k*_IT_. We note that *k*_IT_ could naturally be tuned by evolution.

### An IT-based search can explain the RNA-length-dependence of Cas13 activity

ITs are only useful if *k*_IT_ is small enough that the RNA is mostly able to re-equilibrate between each IT. (We verify the self-consistency of this assumption in Supp. Section S4). Under this assumption, we can sum over *m* to get the total rate of the free binding patch encountering any stretch of RNA along the chain, which will be used to calculate the probability that the dissociation of a binding patch results in an IT rather than in a complete dissociation of the protein. This rate depends on the number of sites upstream and downstream of where the protein is bound, which we denote by *L*_u_ and *L*_d_, respectively:

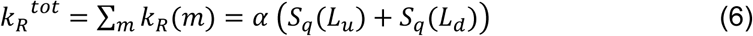

where *α* ∼ *k*_B_*T*/*ηb*^3^ has units of inverse time, *q*=3*l*^2^/2*b*^2^, and we define

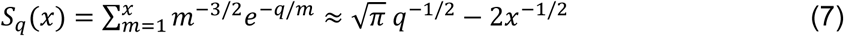

where the approximation is valid in the regime 1 < *q* < *x* (see derivation and approximations for other parameter regimes in Supp. Section S3; see also Supp. Fig. 4). Eqs. 6-7 reveal a key feature of ITs for RBPs, which does not apply to DNA-binding proteins. Recognizing that the total number of ITs taken before dissociating, *n*^IT^, is approximately given by *k*_R_^avg^/*k*_IT_ where *k*_R_^avg^ is the average of *k*_R_^tot^ taken over all positions (see Supp. Section S4), these equations indicate that the number of ITs performed by an RBP is dependent on the length of the RNA to which it is bound: the RBP will perform more ITs on longer RNAs, and fewer ITs on shorter RNAs (Fig. 3F). Or, put another way, an RBP using ITs will dissociate more slowly from longer RNAs, and more quickly from shorter RNAs. As shown in the figure, this effect is particularly apparent for large values of *l*.

**Figure 4:**
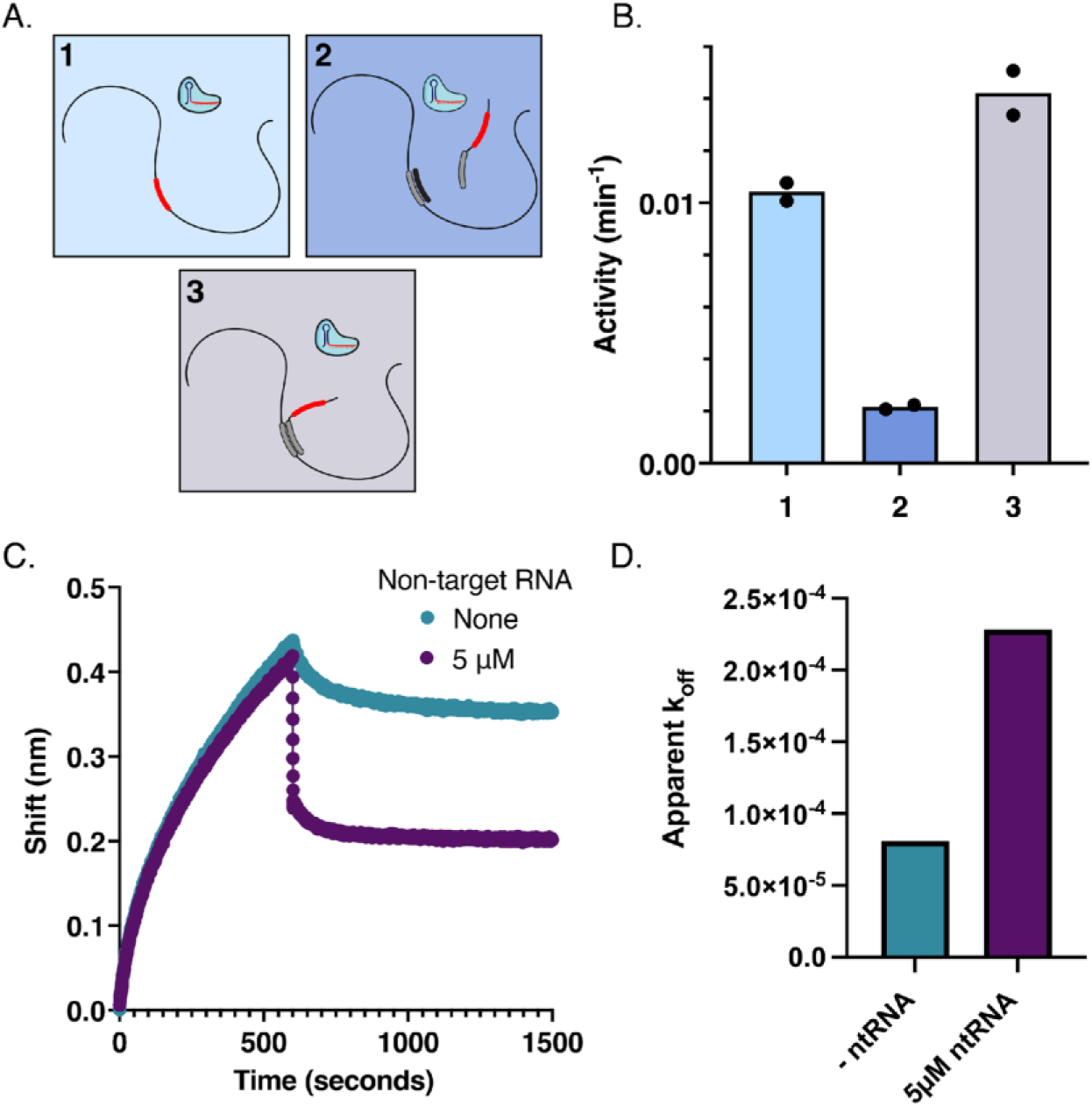
Cas13 uses intersegmental transfers during its search process. **A.** Schematic of an assay to differentiate 1D sliding and intersegmental transfer. In scenario 1, Cas13 targets a protospacer in the middle of a 768 nt target RNA. In scenario 2, Cas13 targets a protospacer on a 71 nt target RNA in the presence of a 768 nt RNA. In scenario 3, the 71 nt target RNA is annealed to the middle of the 768 nt RNA via a complementary region downstream of the protospacer sequence. **B.** Quantification of Cas13 activity for the three scenarios illustrated in F. **C.** Biolayer interferometry data showing that Cas13 dissociates from its target RNA more quickly in the presence of RNA. **D.** Quantification of apparent dissociation rate constants (k_off_) from C.

Moreover, Eqs. 6-7 demonstrate that an RBP performing ITs will experience “edge effects” in qualitative agreement with Cas13’s behavior. When the protein is bound at a certain position, the probability of it performing a successful IT from that position rather than dissociating completely from the RNA is given by (1+*k*_IT_/*k*_R_^tot^)^-1^. As *k*_R_^tot^ is a concave function of position along the RNA, the likelihood of the protein binding to the center of the RNA is higher than the likelihood it binds to the edge of the RNA (Fig. 3G), in agreement with our findings in Fig. 2H. As shown in Fig. 3G, the width of the “edge” grows with the distance *l* between the binding patches on the protein (see Supp. Section S4).

To explore the ability of our RNA-IT model to explain the dependence of Cas13 activity on RNA length, which could not be recapitulated by models of sliding or intracoil diffusion, we consider the dependence of *k*_on_ on *L*_t_. We plot this dependence in Fig. 3H, for *l*/*b*=8. We find that unlike the corresponding plot for a sliding-based search (Fig. 3B) which is concave for all *L*_t_, an IT-based search yields a convex dependence of *k*_on_ on *L*_t_ for small *L*_t_ (Fig. 3H). This prediction shows good agreement with our experimental findings for Cas13 (Fig. 3A). This convexity is present only when *q* ≳1 (i.e. l ≳ b): in this case, for small *L*_t_, the RNA is too short for ITs to transport the protein to an arbitrary segment of the RNA, reducing the efficacy of ITs for short targets.

Similarly, ITs can also explain the effect of non-target RNA sonication on Cas13. The key feature is that when performed on RNAs, ITs lead the protein to dissociate faster from shorter RNA molecules. When Cas13 binds a short RNA at one non-specific binding patch, the probability of a segment of the RNA reaching the second binding patch is small, leading Cas13 to dissociate without performing many ITs. For this reason, higher amounts of sonication of non-target RNAs leads to higher Cas13 activity, as Cas13 wastes less time searching shorter non-target RNAs (Fig. 3I). This prediction is also in good agreement with our experimental results, which displayed a surprising increase in Cas13 activity with greater sonication of non-target RNAs (Fig. 3C). Thus, Fig. 3 demonstrates that Cas13’s responses to changes in the lengths of both target and non-target RNAs can be explained by a target search dominated by ITs, but not by a search dominated by other modes of facilitated diffusion.

### Further experiments confirm that Cas13 employs ITs in its search process

In addition to enabling the protein to transfer between different regions of the same RNA molecule, ITs can also enable transfers between different (nearby) molecules. As searches using 1D sliding do not involve such intermolecular transfers, the ability of the protein to transfer intermolecularly can be used to distinguish searches dominated by sliding from those dominated by ITs. We therefore designed two assays to measure Cas13’s ability to transfer between different RNA molecules.

The first such assay we designed takes its inspiration from Gowers and Halford’s study of the restriction enzyme EcoRV^53^. We tested Cas13’s activity in three distinct scenarios (Fig. 4A): (1) with the protospacer in the middle of a long RNA; (2) with the protospacer in the middle of a short RNA, in the presence of a long non-target RNA; (3) with the protospacer in the middle of a short RNA which has been hybridized to a long non-target RNA. The essential test is in scenario (3), as a search using ITs is expected to find the target at different rates for scenarios 2 & 3, while a search using sliding is expected to find the target in scenarios 2 & 3 at similar rates. The reason for this is that ITs enable the protein to transfer between different nearby RNA chains, while a search using sliding is restricted to the RNA chain to which it initially bound non-specifically. In other words, ensuring the long and short RNAs are spatially co-localized is expected to significantly accelerate a search dominated by ITs, but not a search dominated by sliding. In agreement with our hypothesis that Cas13 uses ITs in its search, we find similar rates of target acquisition for scenarios 1 & 3, and a substantially slower rate for scenario 2 (Fig. 4B, Supp. Fig. 1).

Motivated by the same principles, we designed an additional assay, in which we measured the dissociation rate of Cas13 from the target RNA as a function of the concentration of non-target RNAs in solution. A protein performing a sliding-based search would have the same dissociation rate, regardless of how much non-target RNA was in solution; however, a protein whose searches are dominated by ITs would dissociate much faster in the presence of a high concentration of non-target RNAs. This is because ITs are realized by many microscopic dissociation and reassociation events performed by each binding patch; if a non-target RNA is transiently located near the protein, one of these reassociation events can result in the protein transferring to the non-target RNA. Using biolayer interferometry to directly study binding and dissociation rates, we found that Cas13 binds strongly to its target RNA and, in the absence of non-target RNA, dissociates slowly. However, when non-target RNA is added to the dissociation buffer, the rate of dissociation is greatly increased (Fig. 4C,D). Both this and the previous assay indicate that the mechanism of facilitated diffusion employed by Cas13 is better explained by ITs than by 1D sliding.

### Search time analysis indicates an optimal rate of performing ITs

To understand the evolutionary forces acting on proteins performing ITs, we sought to model how the optimal search would depend on various system parameters. First, we consider a system with no non-target RNAs, using Eq. 1 with *n=n*^IT^+1 and *τ*_t_ ^local^=1/*k*^off^, where *k*^off^=*k*_IT_/(*n*^IT^+1) is the dissociation rate of the protein (see Supp. Section S4). Note that these are not independent, but are both functions of *L*_t_ and of *k*^IT^. In Fig. 5A, we plot the target acquisition rate *k*_on_ as a function of *k*_IT_ for different values of *k*_R_^avg^; Fig. 5B shows a similar plot for different concentrations of target RNA. Although models with independent *k*^off^ and *n*^IT^ can naturally lead to search times being optimized by the protein never dissociating from the target^44^, Fig. 5A-B demonstrate that coupling the two leads the search time to be optimized at a finite value of *k*_IT_. This optimal value is given by

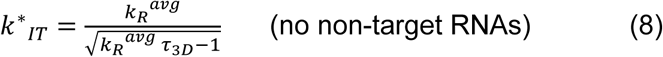

**Figure 5:**
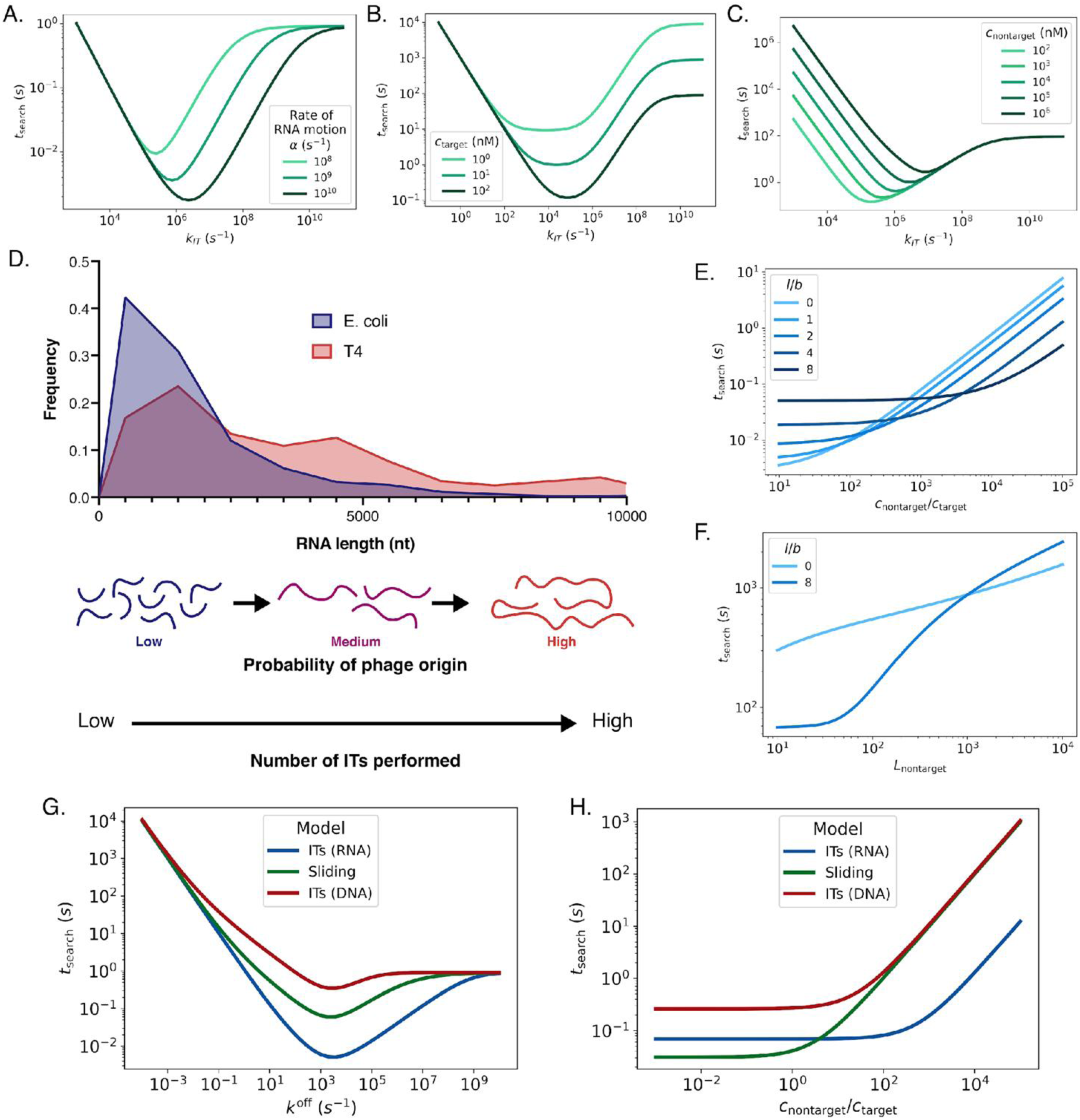
Modeling reveals general advantages of intersegmental transfer in RNA-based search processes. **A.** Curves showing the dependence of search time on the rate of intersegmental transfer, for different rates of overall RNA binding. **B.** Modeling results as in A, but for different concentrations of target RNA in the solution. **C.** Modeling results as in A, but for different concentrations of non-target RNA in the solution. **D.** Histogram of the known RNA length distributions for E. coli and bacteriophage T4. **E.** Curves showing the dependence of search time on the ratio of non-target to target RNA in the system, for different values for the distance between binding domains. **F.** Modeling results as in E, but for different non-target RNA lengths. **G.** Direct comparison of search time for three different modes; ITs searching DNA, sliding, and ITs searching RNA. **H.** Modeling the search time for the same three search strategies as a function of the ratio of non-target to target RNA.

Larger values of *k*_IT_ are suboptimal as they lead to faster dissociation and therefore fewer sites searched per round of binding, while smaller values of *k*_IT_ are suboptimal as they lead each IT to take longer. In the regime where *τ*_3D_<(*k*_R_^avg^)^-1^, *k\**_IT_ is negative and ITs slow down the search for any finite value of *k*_IT_. However, given that *k*_R_^avg^ is of order inverse nanoseconds, this regime is not physically relevant for RNA. We note that the presence of non-target RNAs increases the optimal value of *k*_IT_ compared to Eq. 8, since time that the protein spends searching non-target RNAs before dissociating from them is wasted (Fig. 5C).

The corresponding optimal search time is given by

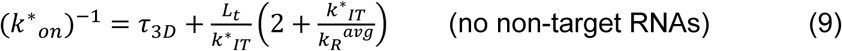

Recognizing that *L*_t_/*k*_IT_ represents the total time the protein spends performing ITs, and that *k*_R_^avg^≫*k\**_IT_, this indicates that, aside from the first 3D search to find the target, the protein optimally spends nearly the same amount of time searching locally through ITs as searching globally through 3D diffusion. Similar results have previously been found for the optimal search using 1D sliding^32^ suggesting that equipartitioning local and global search times may be a feature of facilitated diffusion more broadly.

### Benefits of ITs for a search on RNA

Having found that Cas13 employs ITs in its search, we asked whether such a mechanism may benefit RNA-based search processes more generally. First, Eq. 5 indicates that ITs may be substantially faster on RNA than on DNA, because of the dependence of *k*_R_ on the persistence length. For single-stranded RNA as we consider here, *b*≈2 nm, while for double-stranded DNA *b*≈100 nm^49,52^. For RNA, *k*_R_∼*b*^-3^ as the internal polymer motion is Zimm-like (Eq. 5); for DNA, the motion is more appropriately described by the Rouse model^48^ and the scaling is moderated somewhat^54^ to *k*_R_∼*b*^-2^. Therefore, put together, ITs can be performed 3-5 orders of magnitude faster on RNA than on DNA.

In addition to its effect on the overall search rate, the short persistence length of RNA has other qualitative effects on an IT-based search. In particular, for DNA, an IT-based search will always have *l* ≪ *b* (i.e. nearby binding patches, relative to the persistence length of DNA), while for RNA, *l*≳*b* can be relevant. Indeed, the qualitative features of Cas13’s search time scaling with target RNA length (Fig. 3) can only be understood in the context of large values of *l*/*b* (i.e. a large separation between the binding patches). On its surface, this appears suboptimal, as increased values of *l* lead to fewer ITs performed on each RNA (Fig. 3F) and ultimately to slower searches in the absence of non-target RNA (Fig. 5E; left side of the plot). The key to making sense of this finding is recognizing that an optimal search process must not only find the target molecule quickly, but avoid non-target RNAs as much as possible. When the solution contains many short non-target RNAs, large values of *l* enable the protein to dissociate quickly from these RNAs. As described in Eq. 7, an IT-based search enables the protein to dissociate faster from short RNAs than long ones. As seen in Fig. 3F, especially for *l*≫*b*, the dissociation rate from short RNAs can be orders of magnitude faster than the corresponding rate for long RNAs.

To explore this idea and its potential advantages, we used publicly available datasets to analyze the lengths of all known transcriptional units in two well-characterized systems, E. coli and bacteriophage T4^55,56^. We found that phage RNAs tend to be longer, on average, than bacterial RNAs, and thus infer that a mechanism which prioritizes thoroughly searching long RNAs while avoiding shorter ones will be highly efficient for a phage RNA-targeting system like Cas13, or other RNA-targeting CRISPR systems (Fig. 5D). In this sense, we propose ITs may be acting as a kind of length filter, in order to selectively enrich for longer RNAs likely to be of bacteriophage origin during the search process and thus help narrow the search space.

In Fig. 5E, we consider a solution consisting of long target RNAs (*L*_t_=10^3^) and short non-target RNAs (*L*_i_=40). We plot the search time as a function of the relative concentration of the non-target RNA for proteins with different amounts of separation between the binding patches (i.e. different values of *l*/*b*). While for small amounts of non-target RNA, the search is much faster for proteins with nearby binding patches (i.e. smaller values of *l*/*b*), for high enough non-target RNA concentrations, the search is optimized by proteins with large separations between the binding patches. The reason for this concerns the length-dependent dissociation rate of an RBP performing ITs: even though a protein whose binding patches are farther apart will dissociate faster from all RNAs, it will especially dissociate faster from short RNAs.

Similarly, in Fig. 5F, we plot the search time as a function of the length of non-target RNAs for given values of the non-target RNA concentration and of *l*/*b*. We find that even though large values of *l*/*b* slow down the rate at which the protein can perform each IT, they enable the protein to avoid short non-target RNAs, and thus can speed up a search in which short non-target RNAs are present at high quantities.

Finally, to compare between sliding and ITs more directly for an RNA-based search, in Fig. 5G we plot the search time *t*_search_=(*k*_on_)^-1^ for ITs (blue), as well as for a sliding-based search (green) as a function of the dissociation rate of the protein from the RNA, *k*^off^. For very large values of *k*^off^, the two searches are equivalent to a pure 3D search, and for very small values both are inefficient as the protein gets stuck searching the non-target RNAs. For intermediate dissociation rates, ITs are substantially faster than sliding, given realistic 1D diffusion coefficients of *D*_1D_=10^6^ nm^2^/s (at *D*_1D_=10^9^ nm^2^/s, sliding and ITs are similarly fast). ITs are particularly faster in the presence of non-target RNAs; in their absence, sliding can be faster than ITs (Fig. 5H). The speed of ITs is dependent on the short persistence length of RNA; even in the absence of non-target molecules, our model finds ITs to be a relatively inefficient method of searching DNA (Fig. 5G-H, red).

## DISCUSSION

In this study, we used a feedback loop of biophysical modeling and experimental validation to study how RBPs can quickly find their RNA targets among the vast number of cellular RNAs. While our modeling work is able to consider generic RBPs, our experimental studies focused on a model RBP, Cas13. We established an experimental system that uses Cas13 binding and activation kinetics as a proxy for RNA search speed. We then showed that Cas13 uses facilitated diffusion in its target search process and considered several potential mechanistic models. Multiple orthogonal lines of evidence, including the effect of RNA lengths on Cas13 search time and Cas13’s ability to transfer between different RNA molecules without dissociating, indicate that Cas13 uses a search dominated by intersegmental transfers.

We find that the search process of RBPs differs from that of DNA-binding proteins in three primary ways, all of which lead ITs to be particularly efficient for RNA searches. First, the short persistence length of RNA compared to DNA means that ITs can occur ∼10^4^ times faster on RNA than on DNA. Second, the secondary structures formed by RNA can create obstacles that impede 1D diffusion^27^ which can be circumvented by ITs (Fig. 4A-B). Third, cellular RNAs are heterogeneous in length; a protein which spends equal time on each local search would search short RNAs many times over before ever fully searching a long RNA. While the increased radius of gyration of longer RNAs coupled with intracoil diffusion can partially address this inefficiency, ITs enable a search that is counterintuitively more efficient for long RNAs than for short ones. In particular, ITs enable the protein to naturally (and without any energy consumption) dissociate quickly from short RNAs and spend more time searching longer RNAs. This “length-filtering” effect is strengthened when the two binding patches on the protein are far apart, representing a tradeoff in the position of these patches: when the patches are closer together, the protein searches more sites on each RNA before dissociating; when they are farther apart, the protein searches fewer sites, but in a length-dependent manner, dissociating particularly quickly from shorter RNAs.

For this reason, our results indicate that ITs may be a generically efficient facilitated diffusion mechanism for a broad class of sequence-specific RNA-targeting proteins. Those proteins whose RNA targets are longer than the typical cellular RNA, for instance proteins targeting mRNAs, can utilize the length-filtering effect we have described by placing their binding patches farther apart (yielding qualitatively similar behaviors to our findings on Cas13), while those proteins whose RNA targets are shorter than the typical cellular RNA can place their patches closer together, eschewing length-filtration in favor of faster overall search. In contrast to previous work which considered ITs primarily for multimeric proteins with multiple recognition domains, we propose that ITs may be broadly applicable for proteins with only one recognition domain but multiple non-specific binding patches. While the non-specific binding patches are unable to distinguish the target sequence from non-target themselves (see Supp. Section S4), they facilitate faster binding between the specific binding domain and the target by enabling the protein to perform ITs. For example, Cas13 has only one RNA-recognition domain which can distinguish the specific target sequence from other RNAs, but positively charged patches on its surface can serve as separate locations for non-specific RNA association (Supp. Fig. 5). In fact, RBPs have been found to be highly enriched for such positively charged patches, and roughly 45% of eukaryotic RBPs contain multiple experimentally validated RNA-binding domains^57,58,59^. The search behavior of several RBPs appears to also be consistent with ITs^27–29,41^, though ITs have rarely been considered due to their relative inefficiency on DNA compared to sliding^45,60^.

While we have treated them separately in this work, ITs have a close connection to intracoil diffusion. In the limit of fast polymer diffusion compared to protein diffusion, an intracoil diffusion process may be qualitatively similar to ITs with *q*=0. The model we use for intracoil diffusion in this work is appropriate in the opposite limit. Developing a comprehensive theory of the intracoil diffusion process across different regimes of polymer diffusion rates may be a fruitful subject for future work^61^. Intracoil diffusion, as well as 1D sliding, may indeed play a substantial role in Cas13’s search process; our model does not rule out these mechanisms entirely but rather shows that these alone cannot explain Cas13’s search behavior. Intracoil diffusion occurs as part of the 3D diffusion process, and it is possible that a mixture of longer-distance transfers and local 1D searches may lead to more efficient search. Future work will identify what role, if any, is played by other search modalities beyond ITs in Cas13’s search process.

In an effort to isolate the RNA search process and minimize confounding factors, we performed all assays in highly controlled environments using carefully designed RNAs.

Therefore, this work neglects many features of the cellular environment that may influence the search process *in vivo*, including different secondary structures, the effects of other RNA-binding proteins, and molecular crowding. These effects will be the subject of future work. Our IT model will also be expanded upon, to include effects such as asymmetry between binding patches, the presence of more than 2 patches, or extended binding regions.

Though we have framed the RNA search problem as it pertains to Cas13 operating in its native prokaryotic context, Cas13 has also been broadly utilized in eukaryotic cells for RNA knockdown, base editing, and RNA imaging. In these contexts, the search problem is even greater; eukaryotic cells are on average around 1,000 times larger by volume and contain hundreds of times as many RNAs spread across multiple cellular compartments ^4,62,63^. Thus, improved understanding of the RNA search process is critical for furthering the biotechnological applications of Cas13, as it may lead to improvements in RNA targeting efficiency via intelligent crRNA design, protein engineering, or other approaches.

## ACKNOWLEDGMENTS

We thank Long Nguyen and Soonwoo Hong for protocols and technical assistance related to protein purification, and Owen Abad for helpful discussions and preliminary experimental planning. We thank Venu Vandavasi and the Princeton Biophysics Core Facility for access to and training on the Octet RED96e instrument. We thank Alexander Grosberg for discussions regarding the scaling of intracoil diffusion with target length. We thank Ned Wingreen and the members of the Myhrvold and te Velthuis labs for helpful discussions.

This work was supported by the Branco Weiss Fellowship–Society in Science, administered by the ETH Zürich (O.K.), the Peter B. Lewis ’55 Lewis-Sigler Institute/Genomics Fund (O.K.), the National Institutes of Health (R01AI182281, C.M.; R01AI177487, A.J.W.t.V.; and DP2AI175474, A.J.W.t.V.; T32GM007388, B.B.L; T32GM148739, B.B.L and E.G.), and the Princeton Catalysis Initiative (C.M.). Research reported in this publication was supported by the National Center For Advancing Translational Sciences of the National Institutes of Health under Award Number UM1TR004789. The content is solely the responsibility of the authors and does not necessarily represent the official views of the National Institutes of Health.

## METHODS

### Target RNA design

To control for effects of RNA secondary structure, we designed target RNAs to include minimal structure, as described previously in Ref.^37^. For most experiments, crRNA 4 and its corresponding protospacer were used, as that crRNA showed high activity in our previous studies, and was compiled from the ADAPT dataset which has a sequence composition representative of viral diversity^36^. The data labeled “protospacer 2” in Fig. 2D represent results using crRNA 5 from the same paper; in other experiments, RNAs including the protospacer corresponding to crRNA 5 were used as non-target RNA. RNAs of different lengths were created by concatenating the protospacer with RNA sequences compiled out of a set of 64 16-nt-long DNA sequences with minimal internal structure from Shortreed *et al.*^64^. The protospacer was placed in the middle of the sequence, aside from the experiments described in Fig. 2G-H which included 5 equally-spaced identical protospacers. NUPACK^65^ was used to predict the structure of the resulting sequences and ensure minimal secondary structure. All sequences used are shown in Supp. Table 1.

### RNA preparation

Templates for target RNAs for use in Cas13 assays were purchased from IDT (kinetics experiments and BLI) or Ansa Biotechnologies (DMS experiments) as DNA. Templates containing a double-stranded T7 promoter sequence were transcribed to RNA using homemade T7 polymerase mix. For a 20 μL IVT reaction, 1.5 μL enzyme master mix, 10 μL 2× IVT reaction mix, 1 μL 0.1 M DTT, DNA template, and nuclease-free water were combined. The 2× IVT reaction mix contained 20 mM rNTP mix, 80 mM Tris-HCl, 12 mM MgCl₂, 20 mM DTT, and 2 mM spermidine. Reactions were carried out at 37 °C for 16 hours followed by DNAse I XT template digestion and subsequent purification using 1.8× SPRI beads and the addition of 1.6× isopropanol for RNAs of length < 100 nt, and were eluted into 20 μL of nuclease-free water.

RNAs were quantified using a NanoDrop One (Thermo Fisher Scientific) or Biotek Take3Trio (Agilent) and stored in nuclease-free water at −80 °C.

When pre-annealing RNAs or DNA-RNA hybrids, equimolar amounts of the two species (or a 2:1 ratio of DNA to RNA in certain cases) were mixed in a 60 mM KCl annealing buffer, then annealed in a thermocycler by heating to 85 °C and gradually cooling to 10 °C at 0.1 °C/second, before final cooling to 4 °C.

### Cas13 activity assays

Plate reader detection assays were performed by mixing target RNA at a ratio of 10% v/v with 90% Cas13 detection mix. The detection mix consisted of 1× SHINE buffer (20 mM HEPES pH 8.0, 60 mM KCl and 3.5% PEG-8000 in nuclease-free water), supplemented with 45 nM purified homemade LwaCas13a, 1 U per μl murine RNAse Inhibitor (New England Biolabs), 62.5 nM fluorescent reporter (/5FAM/rUrUrUrUrUrU/IABkFQ/; IDT), 22.5 nM processed crRNA (IDT) and 14 mM magnesium acetate. Reactions (15 μL each) were then loaded in technical duplicate onto a 384 well clear-bottom Greiner plate. The plate was then loaded onto a Biotek Cytation5 or Synergy H1 plate reader, and fluorescence intensity was measured every 5 minutes for 3 hours at 37 °C.

Highly multiplexed chip-based assays were performed using a BioMark HD system with a Standard Biotools genotyping IFC (96.96 format). The assay mix (7% of reaction volume) contained 1× Assay Loading Buffer (Standard Biotools), 293 nM LwaCas13a, and 1× SHINE buffer (see above section). The sample mix (93% of reaction volume) contained 1.07 U/μL murine RNAse Inhibitor (New England Biolabs), 15 mM MgOAc, 110 nM FAM reporter, 1× ROX reference dye (Standard Biotools), 1× GE buffer (Standard Biotools), 1× SHINE buffer, and target RNA (variable concentration).

Sample volumes of 6 μl and assay volumes of 6 μl, in addition to control line fluid, were loaded onto the IFC chip (Standard Biotools). Chips were then placed into the IFC Controller HX and primed, loaded and mixed using the Prime and Load Mix 96.96 GE scripts (Standard Biotools). After mixing, chips (with two technical replicates per condition) were run on BioMark HD at 37 °C for 8 h, with measurements taken in the FAM and ROX channels every 5 min. Normalized and background-subtracted fluorescence for a given time point was calculated as (FAM − FAM background)/(ROX − ROX background).

For the strand displacement assay, reaction conditions were the same as standard Cas13 detection assays, with the following adjustments: the strand-displacement reporter concentration was set to 20 nM, the dCas13 concentration (in the +dCas13 condition) was set to 9 nM, and the crRNA concentration was set to 4.5 nM.

### Protein purification

For LwaCas13a: a plasmid containing the human codon-optimized LwaCas13a sequence (Addgene ID: 90097) was transformed into Rosetta2(DE3) Singles™ Competent Cells according to the manufacturer’s protocol. Single colonies were inoculated into 30 mL LB broth and incubated at 37 °C overnight. The following day, overnight cultures were inoculated into 2 L Terrific Broth and incubated at 37 °C with shaking at 250 rpm until reaching an OD of 0.6. Cultures were then placed on ice and induced with IPTG at a final concentration of 0.5 mM and cultured overnight at 18 °C with shaking.

The cells were then pelleted by centrifugation (4000 ×g, 10 min) and resuspended in lysis buffer (20 mM Tris-HCl pH 7.5, 500 mM NaCl, 20 mM imidazole, 1 mM DTT) supplemented with protease inhibitors (Complete Ultra EDTA-free tablets) and lysozyme (500 μg). Cells were then lysed by sonication (10 s on/10 s off for 20 min with 50% power) and the lysate subsequently clarified by centrifugation (20,000 ×g, 1 hr, 4 °C). The supernatant was passed through a 0.2 micron filter before being injected onto the Biorad NGS FPLC system connected to a Ni-NTA column and purified using a wash buffer (20 mM Tris-HCl pH 7.5, 500 mM NaCl, 1 mM DTT) and an elution buffer (20 mM Tris-HCl pH 7.5, 500 mM NaCl, 300 mM imidazole, 1 mM DTT). Eluted fractions were pooled and mixed with with 300 μL of 1 mg/mL SUMO protease then dialyzed against SUMO cleavage buffer (30 mM Tris HCl pH 8.0, 500 mM NaCl 1 mM DTT, 0.15% Igepal (NP-40)) overnight. The protein was concentrated to 50% of its original volume using an Amicon 50 kDa MWCO spin column then mixed at a 1:1 ratio with cation exchange buffer (20 mM HEPES pH 7.5, 250 mM NaCl, 1 mM DTT, 5% glycerol). The protein was then further purified by injecting it onto a 5 mL HiTrap SP HP cation exchange column and eluted over a salt gradient of 250 mM to 2 M NaCl in elution buffer (20 mM HEPES pH 7.5, 2 M NaCl, 1 mM DTT, 5% glycerol).

Protein purity was checked on a Coomassie-stained PAGE gel, after which fractions were pooled. If purity was not sufficient, the sample was subjected to an additional purification step consisting of size exclusion chromatography over a Superdex 200 Increase 10/300 GL column. The protein was then concentrated and exchanged into storage buffer (600 mM NaCl, 50 mM Tris-HCl pH 7.5, 5% glycerol, 2 mM DTT) again using a 50 kDa MWCO spin column and subsequently aliquoted and flash-frozen in liquid nitrogen. Aliquots were stored at -80 °C until use.

For T7 RNA polymerase, RNAse inhibitor, and IPPase: The T7 RNAP variant (G47A+884G) coding sequence was cloned into a pET-derived N-terminal 6× His-SUMO expression vector. The RNase inhibitor expression construct was generated in-house, and inorganic pyrophosphatase (iPPase) was expressed from a gift from Sebastian Maerkl and Takuya Ueda (Addgene plasmid #124137).

RNA Bacterial expression plasmids were transformed into Rosetta 2(DE3)pLysS Singles competent cells (Millipore Sigma, Cat. #71401). Individual colonies were picked into 50 mL Luria Broth (Fisher Scientific, Cat. #BP9723-2) containing the appropriate antibiotic(s) and grown overnight at 37°C with shaking. Overnight cultures were used to inoculate 4 L Terrific Broth (RPI, Cat. #T15000-10000.0), and cultures were grown at 37 °C until the OD600 reached 0.6–0.8.

Cultures were cooled rapidly on ice for 10–15 min, induced with 1 mM IPTG (Gold Biotechnology, Cat. #I2481C100), and incubated overnight at 18°C. Cells were harvested by centrifugation at 4,000 ×g for 15 min. Cell pellets from 4 L culture were resuspended in 150 mL lysis buffer (50 mM Tris-HCl pH 7.9, 5% glycerol, 500 mM NaCl, 20 mM imidazole, and 1 mM TCEP). Lysozyme (0.5 mg/mL; Astatech, Cat. #AT25331 5G), DNase I (0.01 mg/mL; Worthington, Cat. #LS002145), and PMSF (0.5 mM; Tokyo Chemical Industry, Cat. #B1062) were added to the lysis buffer. Cells were lysed by sonication on ice. Lysates were clarified by centrifugation at 186,000 ×g for 30 min using a Beckman Coulter Type 45 Ti rotor, and the supernatant was passed through a 0.45 μm filter. Clarified lysate was loaded onto a prepacked Ni-NTA affinity column (EconoFit Nuvia IMAC column, Bio-Rad, Cat. #12009287) using a Bio-Rad NGC Quest Plus FPLC system. Bound protein was washed with lysis buffer and eluted with elution buffer (50 mM Tris-HCl pH 7.9, 5% glycerol, 500 mM NaCl, 400 mM imidazole, and 1 mM TCEP). Fractions containing the target protein were identified by SDS-PAGE and pooled.

For T7 RNAP, pooled Ni-NTA eluate was digested with SUMO protease to remove the N-terminal 6× His-SUMO tag. Briefly, eluate was combined with 2 mg SUMO protease in dialysis tubing (Ward’s Science, Cat. #470163-414) and dialyzed overnight at 4°C with rotation against SUMO digestion buffer (30 mM Tris-HCl pH 8.0, 500 mM NaCl, 1 mM DTT, and 0.15% Igepal/NP-40,). Following digestion, the protein was buffer-exchanged into buffer A (50 mM Tris-HCl pH 7.9, 5% glycerol, 200 mM NaCl, and 1 mM TCEP), using 50 kDa MWCO centrifugal filters (Millipore Sigma, Cat. #UFC905024). The buffer-exchanged sample was applied to a 5 mL HiTrap Heparin HP column (Cytiva, Cat. #17040701), and T7 RNAP was eluted using a linear gradient from buffer A to buffer B (50 mM Tris-HCl pH 7.9, 5% glycerol, 2 M NaCl, and 1 mM TCEP). Fractions containing purified T7 RNAP were pooled.

Purified T7 RNAP, RNase inhibitor, and iPPase were buffer-exchanged and concentrated using 50 kDa MWCO centrifugal filters into storage buffer (50 mM Tris-HCl pH 7.9, 5% glycerol, 100 mM NaCl, 0.1% Triton X-100, 1 mM EDTA, and 1 mM DTT). Proteins were individually aliquoted and stored at 80 °C. In-house IVT enzyme master mix was prepared by combining purified T7 RNAP, RNase inhibitor, and iPPase at final concentrations of 2.67 μg/μL, 2.67 μg/μL, and 0.27 μg/μL, respectively, in a buffer containing 50 mM Tris-HCl pH 7.9, 100 mM NaCl, 50% glycerol, 0.1% Triton X-100, 1 mM EDTA, and 1 mM DTT. The enzyme master mix was then stored at -20 °C.

### DMS footprinting assay

The Cas13 binary complex, or crRNA alone, was prepared at a final concentration of 1μM in 1× binding buffer as described above (see “EMSA” section). This complex was then mixed with target RNA in 1× binding buffer at a ratio of 1:4 to ensure excess unbound target was present, and allowed to complex for 15 minutes at 37 °C. Each 10 μL sample was added to 35 μL of refolding buffer (0.38 M sodium cacodylate pH 7.2, 14 mM MgCl_2_) then treated with 7 μL of DMS solution (15% v/v dimethyl sulfate, 85% v/v ethanol) or ethanol-only control at 37 °C. Reactions were quenched by the addition of 50 μL of beta-mercaptoethanol. Samples were cleaned up in two steps; first, they were subjected to column-based purification (Zymo RNA Clean and Concentrator 5), then purified by 5× SPRI beads. Samples were eluted into nuclease free water, quantified by NanoDrop One, and stored for downstream library preparation.

For each sample, 200 ng of RNA was dephosphorylated using Quick CIP (NEB) according to the manufacturer’s instructions, before halting reactions by heat-inactivation at 85°C. Next, a single phosphate group was restored to the 5’ ends of the entire sample using T4 polynucleotide kinase (NEB). The reaction was allowed to proceed at 37 °C overnight, before heat-inactivation at 65 °C for 20 minutes. Each sample was once again purified using 1× SPRI beads and eluted into 10 μL of nuclease-free water. The eluted RNA was then circularized using RNA Ligase I (NEB), with the addition of 20 units of murine RNAse inhibitor. Reactions were allowed to proceed overnight at 16 °C, before heat-inactivation at 65 °C for 15 minutes. First-strand cDNA was synthesized from the circular RNA using UltraMarathonRT (RNAConnect), with the addition of High Boost. The resulting cDNA was then used as the template for PCR amplification using Q5 DNA polymerase (NEB) and primers containing 5’ sequencing barcodes. The PCR products were purified and size-selected using SPRI beads, eluted into nuclease-free water, then pooled for next-generation sequencing.

### DMS assay next-generation sequencing analysis

The reads were first demultiplexed using Barcode Splitter, and subsequently the adapters were trimmed using cutadapt. The reads were then split into the regions 5’ and 3’ to the ligation site in order to remove gaps and nontemplated bases inserted by T7 polymerase or uMRT. Cutadapt was used to remove barcodes (--cut) and to trim sequences to the desired length (--length). The two sections were then separately aligned to their respective sections of the template using Bowtie2 with settings --local, -N 1, -L 18. Reads with MAPQ (mapping quality) score <25 were discarded. A custom script was used to analyze the per-nucleotide mutation rate for each sample, using the rate of mutagenesis per A nucleotide (excluding primer-binding regions) to generate an overall mutation rate per protospacer.

### Biolayer interferometry

Biotinylated DNA adapters were pre-annealed to target RNAs containing a 3’ complementary region, and the duplexes were then immobilized on streptavidin biosensors (FortéBio). The sensors were then equilibrated in glycerol-free protein storage buffer (50 mM Tris-HCl pH 7.5, 600 mM NaCl, 2 mM DTT) before submerging into 100 nM Cas13-crRNA complex (prepared as previously described) and allowed to associate for 10 minutes at 37 °C. Sensors were then moved to fresh storage buffer either containing or lacking non-target RNA, and allowed to dissociate for 20 minutes at 37 °C. All data collection and processing were performed on an Octet RED96e biolayer interferometry system using Octet Data Analysis HT software version 11.1.3.50.

### Activity fits and plotting

Activity scores were calculated by fitting fluorescence data to pseudo-first order kinetic curves (one-phase association, GraphPad Prism version 10.6), then defining “activity” as the rate constant (k, with units of inverse time). Separate curves were fit for each technical replicate and used to calculate descriptive statistics per condition. Curves and corresponding activity scores were plotted in Prism.

### RNA length distribution analysis

Bacteriophage T4 transcription units were determined using promoter and terminator annotations from Table 1 of Ref. ^56^. Transcript lengths were determined by calculating the distance between the genomic location of each promoter and the nearest terminator, accounting for the direction (+ strand or - strand) of the promoter and terminator. All promoters and terminators were assumed to be unidirectional based on their annotations in the table. *E. coli* transcription units were taken from Supplementary Table 12 of Ref. ^55^. Transcription unit lengths were determined by subtracting the “Start” and “End” columns of the table.

