## Supplementary Information for "Intersegmental transfers drive target search in an RNA-targeting CRISPR system"

### Section S1: Search time model

Eq. (1) has been derived in several different methods, as referenced in the main text. Below, we present another derivation, highlighting that Eq. (1) is generically applicable regardless of the mechanism of local search. Our derivation is motivated by Appendix A of Ref. [1].

We consider the RNA to be composed of $L$ sites, with the target at position $i^*$. We set up the differential equations governing the rate of transitions into and out of each site:

$$\frac{du_i}{dt} = \frac{1}{L\tau_{\text{3D}}} e^{-t/\tau_{\text{3D}}} + \sum_j k_{ji} u_j - \sum_j k_{ij} u_i - k_{\text{off}} u_i + \frac{k_{\text{off}}}{L\tau_{\text{3D}}} \int_0^t dt' e^{-(t-t')/\tau_{\text{3D}}} u_j(t') \quad \text{(S1)}$$

The terms can be understood as follows:

We denote by $u_i(t)$ the probability that a searching protein will be at site $i$ at time $t$. All terms $u_i$ in the equation above are implicitly functions of $t$. In a system with $N > 1$ searching proteins, we assume that the proteins are sufficiently dilute that they do not interact with one another.

The first term is the rate of the first association to the RNA strand, which we assume occurs with equal probability at each of the $L$ sites. This exponentially distributed rate has a mean of $\tau_{\text{3D}}$ and ultimately accounts for the first $\tau_{\text{3D}}$ term in Eq. (1).

The second and third terms account for direct transfers between sites, meaning motion along the RNA that occurs without fully dissociating from the RNA. $k_{ij}$ is the rate of direct transitions from site $i$ to site $j$, which we leave here in a completely general form. In our work, this could take the form of 1D sliding, intracoil diffusion, or intersegmental transfers.

The final two terms account for dissociations of the protein from the RNA. The protein dissociates at a site-independent rate $k_{\text{off}}$. When the protein fully dissociates, it associates at a rate $1/\tau_{\text{3D}}$. We assume that reassociation occurs at each site with equal probability.

We denote the target site as $i^*$, and treat it as a sink—once the protein reaches the target site it doesn't leave—such that $k_{i^*j} = 0$ for all $j$, and $k_{\text{off}} u_{i^*} = 0$.

To make progress, we recognize that

$$\frac{du_{i^*}}{dt} = \frac{1}{\tau_{\text{3D}}} e^{-t/\tau_{\text{3D}}} - \sum_{i \neq i^*} \frac{du_i}{dt} - k_{\text{off}} \sum_{i \neq i^*} u_i + \frac{k_{\text{off}}}{\tau_{\text{3D}}} \int_0^t dt' e^{-(t-t')/\tau_{\text{3D}}} \sum_{i \neq i^*} u_i(t') \quad \text{(S2)}$$

Marvelously, all local transfers (not mediated by complete dissociation of the protein from the RNA) cancel out in Eq. (S2), such that our result will be independent of the method of local search.

Eq. (1) describes the mean search time, which can be derived from Eq. (S1) as

$$\begin{aligned} \tau_{\text{search}} &= \int_0^\infty dt\, t \frac{du_{i^*}}{dt} \\ &= \tau_{\text{3D}} - \sum_{i \neq i^*} \int_0^\infty dt\, t \frac{du_i}{dt} - k_{\text{off}} \sum_{i \neq i^*} \int_0^\infty dt\, t\, u_i \\ &\quad + \frac{k_{\text{off}}}{\tau_{\text{3D}}} \int_0^\infty dt\, t \int_0^t dt' e^{-(t-t')/\tau_{\text{3D}}} \sum_{i \neq i^*} u_i(t') \end{aligned} \quad \text{(S3)}$$

The first integral can be addressed by integration by parts; the boundary terms vanish. The last integral can be addressed by flipping the order of integration:

$$\frac{1}{\tau_{3D}} \int_0^\infty dt\ t \int_0^t dt' e^{-(t-t')/\tau_{3D}} u_i(t') = \frac{1}{\tau_{3D}} \int_0^\infty dt' u_i(t') \int_{t'}^\infty t\ dt\ e^{-(t-t')/\tau_{3D}}$$

We then change variables on the second integral to $t'' = t - t'$, yielding

$$\frac{1}{\tau_{3D}} \int_0^\infty dt' u_i(t') \int_0^\infty (t' + t'') dt'' e^{-t''/\tau_{3D}} = \int_0^\infty dt' t' u_i(t') + \tau_{3D} \int_0^\infty dt' u_i(t')$$

Thus, our equation for the search time becomes

$$\tau_{\text{search}} = \tau_{3D} + \left( \sum_{i \neq i^*} \int_0^\infty dt\ u_i(t) \right) (1 + \tau_{3D} k_{\text{off}}) \tag{S4}$$

The term in the large parentheses, $(\sum \int u_i dt)$, represents the typical total time the protein spends non-specifically bound to the RNA. It contributes to the total search time in two ways. First, it is the total time the protein spends performing local searches. Second, when multiplied by $k_{\text{off}}$ it represents the typical number of dissociation events required before the target site is found. Each of these dissociation events results in a global (3D) search that takes a typical time $\tau_{3D}$. By identifying $L/n$ as the typical number of local searches performed, we arrive at Eq. (1).

### Section S2: Effect of RNA length on search time

In the absence of non-target RNA, the rate of non-specific association between the protein and the RNA is given by Smoluchowski's equation [2]. Modeling both as spheres, the rate at which a single protein non-specifically binds a target is given by $k_t^{3D} = 4\pi c_t D_t r_t$ where $c_t$ is the concentration of target RNAs, $D_t$ is the sum of the diffusion coefficients of the protein and target RNA, and $r_t$ is the sum of the radius of gyration of the target RNA and the size of the RNA-binding domain of the protein. Because the RNA radius of gyration is typically much larger than the RNA-binding domain of the protein, we neglect the latter here for convenience.

The RNA radius of gyration is proportional to $L_t^v$ with $v < 1$ being the Flory exponent ($v = \frac{1}{2}$ for a random walk). Similarly, $D_t$ is the sum of the diffusion coefficients of the RNA ($D_t^{\text{RNA}}$) and of the protein ($D^{\text{prot}}$). $D^{\text{prot}}$ is a constant, while $D_t^{\text{RNA}}$ scales inversely with the hydrodynamic radius of the RNA (which is very close to its radius of gyration [3]). The dependence of $k_t^{3D}$ on RNA length therefore has two regimes. Letting $r_t = R\ L_t^v$ and $D_t^{\text{RNA}} = d/L_t^v$:

$$k_t^{3D} = 4\pi c_t R L_t^v (D^{\text{prot}} + d\ L_t^{-v}) \tag{S5}$$

Letting $L_t^* = (d/D^{\text{prot}})^{1/v}$, when $L_t \ll L_t^*$, $k_t^{3D}$ is independent of $L_t$, and when $L_t \gg L_t^*$, $k_t^{3D}$ scales as $L_t^v$. In the main text, we make the assumption that $L_t \gg L_t^*$ for simplicity, but as described below, our results (in particular that our experimental findings are inconsistent with pure 3D diffusion, intracoil diffusion, and 1D sliding) are robust to this assumption.

We will now estimate the numerical values of the parameters. The 3D diffusion coefficients of proteins are typically around $10^5$–$10^8$ nm$^2$/s [4, 5, 6]. To our knowledge, the diffusion coefficient of Cas13 in its Apo form has not been measured, but that of Cas9 has been measured to be approximately $10^7$ nm$^2$/s [7]. The coefficient relating $D_t$ to $1/r_t$ can be derived from the Stokes-Einstein equation and is approximately between $10^7$ and $10^8$ nm$^3$/s in the cell [3, 8]. Equating $L_t$ with the number of nucleotides in the RNA chain, the dependence of $r_t$ on $L_t$ has been estimated through analysis of experimentally determined RNA structures to be $r_t \approx 0.55\ L_t^{1/3}$ nm for folded RNA [9] and through analysis of simulations to be $r_t \approx 0.5\ L_t^{0.38}$ nm for unstructured RNA [3]. These estimates place $L_t^*$ roughly in the range of 100–1000 nts. For an RNA concentration of 1 $\mu$m$^{-3}$, these estimates place $k_t^{\mathrm{3D}}$ on the order of 1/s. A more precise estimate would also include a correction factor accounting for that not every interaction between the protein and RNA is at the precise orientation to lead to binding, thereby decreasing $k_t^{\mathrm{3D}}$ [10].

In the remainder of this section we will derive estimates for the scaling behavior of the search time with $L_t$ for various search strategies.

**Pure 3D search.** In a "pure 3D search", defined here as a search for which $n = 1$, the protein diffuses in 3D, binds to a single spot on the RNA, and if it is not the target, it dissociates and repeats the search. For a pure 3D search, the search time is approximately given by $L_t/k_t^{\mathrm{3D}}$ (neglecting the dwell time of the protein on the RNA in between each 3D diffusion event). This search time monotonically increases with $L_t$ for all values of $L_t$:

$$(k_{\mathrm{on}})^{-1} = L_t \tau_{\mathrm{3D}} = L_t (D^{\mathrm{prot}} L_t^v + d)^{-1} \frac{1}{4\pi c_t R} \left(1 + \tau_d \sum_i k_i^{\mathrm{3D}}\right) \quad \text{(pure 3D diffusion)} \quad \text{(S6)}$$

For $L_t \ll L_t^*$, $\tau_{\mathrm{search}} \sim L_t$ and for $L_t \gg L_t^*$, $\tau_{\mathrm{search}} \sim L_t^{1-v}$.

As noted above, in the absence of non-target RNA and for $c_t = 1\ \mu\mathrm{m}^{-3}$, $\tau_{\mathrm{3D}}$ is approximately 1 s times a correction factor accounting for orientational constraints. Setting this correction factor to 0.1 and considering $L_t = 10^3$ as a rough estimate, Eq. (S6) gives a target acquisition rate of $10^4$ s. To compare to experimental results we consider the second-order rate constant $k_{\mathrm{on}}/c_t$, which is thereby estimated to be $6 \times 10^4\ \mathrm{M}^{-1}\mathrm{s}^{-1}$. This is in good agreement with measurements of *in vivo RNA-RNA* interactions (in the absence of proteins) which have an association rate of $9 \times 10^4\ \mathrm{M}^{-1}\mathrm{s}^{-1}$ [11].

**Intracoil diffusion.** Intracoil diffusion also relies only on 3D diffusion, but because it has $n > 1$ (and is therefore an example of facilitated diffusion as we define it here) we distinguish between intracoil diffusion and a pure 3D search. For intracoil diffusion, $n = aL/r$ where $a$ is the length of each site of the RNA [4]. For notational simplicity, we let $r_t = RL_t^v$ and define a dimensionless parameter $R' = R/a$. We assume that the time spent diffusing intramolecularly is negligible, and further assume a uniform dwell time $\tau_d$ for each non-specific binding event. In this model, a search dominated by intracoil diffusion has:

$$(k_{\mathrm{on}})^{-1} = \frac{1 + R' L_t^v}{4\pi c_t R (D^{\mathrm{prot}} L_t^v + d)} \left(1 + \frac{\tau_d}{R'} \sum_i k_i^{\mathrm{3D}} L_i^{1-v}\right) + L_t \tau_d \quad \text{(intracoil diffusion)} \quad \text{(S7)}$$

For realistic RNA molecules, $L_t \gg 1$ and $R' \gg L_t^{-v}$ (since as argued above, $R \approx 0.5$ nm if $a = 0.33$ nm or 1 nt) and the equation becomes

$$(k_{\mathrm{on}})^{-1} = \frac{1}{4\pi c_t a (D^{\mathrm{prot}} + d\ L_t^{-v})} (1 + \tau_d \sum_i k_i^{\mathrm{3D}} n_i) + L_t \tau_d \quad \text{(intracoil diffusion)}$$

If $\tau_d$ is negligible, for $L_t \ll L_t^*$, $\tau_{\mathrm{search}} \sim (L_t)^v$ and for $L_t \gg L_t^*$, $\tau_{\mathrm{search}} \sim (L_t)^0$. Overall, the search time increases monotonically with target length.

**1D sliding.** A sliding based search has the property that $\tau_t^{\mathrm{local}} = n^2/2D_{\mathrm{1D}}$ with $D_{\mathrm{1D}}$ a 1-dimensional diffusion constant. Unlike intracoil diffusion, $n$ is independent of $L_t$. This yields the following search time:

$$(k_{\mathrm{on}})^{-1} = L_t^{-v} (n + L_t) \frac{1 + \frac{n^2}{2D_{\mathrm{1D}}} \sum_i k_i^{\mathrm{3D}}}{4\pi c_t R (D^{\mathrm{prot}} + d\ L_t^{-v}) n} + \frac{n}{2D_{\mathrm{1D}}} L_t \quad \text{(1D sliding)} \quad \text{(S8)}$$

The first term of this equation contains the non-monotonicity in $L_t$. Consider $L_t \gg L_t^*$: when $n \gg L_t$, it decreases with $L_t$ as $L_t^{-v}$; and when $n \ll L_t$, it increases with $L_t$ as $L_t^{1-v}$. (For $L_t \ll L_t^*$, it is independent of $L_t$ when $n \gg L_t$ and increases with $L_t$ as $L_t$ when $n \ll L_t$). However, this non-monotonicity does not match the non-monotonicity seen in the experiments, as discussed in the main text. Making the assumption $L_t \gg L_t^*$ captures the essential features of the non-monotonicity and leads to Eq. (3) in the main text.

**Modeling sonication.** We model non-target RNA sonication by introducing a parameter $s$. $s = 1$ implies no sonication, meaning that the RNA of length $L_n$ is at concentration $c_n$. Sonication by an amount $s$ represents the RNA being cleaved such that it is of length $L_n/s$ at concentration $sc_n$. In Fig. 3D,I, the x-axis represents $s$.

**Changes to *trans*-cleavage rate.** A potential—ultimately incorrect—explanation for our initial results showing that Cas13 activity decreases with the presence of non-target RNAs (Fig. 1D) is that these non-target RNAs compete with fluorescent reporters for *trans*-cleavage. Or, put another way, that the changes to Cas13 activity we see experimentally are dominated by changes to its *trans*-cleavage behavior rather than by changes to its search rate. However, this hypothesis is inconsistent with our subsequent findings of the non-monotonicity of Cas13 activity with target RNA length (Fig. 3A), as well as with our findings that Cas13 activity increases upon target (Supp. Fig. 3) and non-target (Fig. 3C) RNA sonication. When modulating the RNA target length, we observe large changes in Cas13 activity in agreement with our proposed IT model, despite the lack of additional competition for reporter RNAs; in the case of the sonication experiments, the more highly-sonicated samples exhibit higher Cas13 activity, despite actually containing a higher concentration of non-target RNAs and thus more reporter competition.

### Section S3: The number of IT steps taken before dissociating

In this section we describe how to estimate the sum

$$S_q(x) = \sum_{m=1}^{x} m^{-3/2} e^{-q/m}$$

We will use an Euler-Maclaurin expansion to approximate the sum by integrals. We consider this sum in different regimes of $q$.

For $q < 1$, we expand the exponential as a Taylor series, resulting in

$$\begin{aligned} S_q(x) &= \sum_{m=1}^{x} m^{-3/2} \left(1 - \frac{q}{m} + \frac{q^2}{2m^2} + O(q^3)\right) \qquad (q < 1 < x) \\ &= \sum_{m=1}^{x} m^{-3/2} - q \sum_{m=1}^{x} m^{-5/2} + \frac{q^2}{2} \sum_{m=1}^{x} m^{-7/2} + O(q^3) \end{aligned}$$

Each term in this series is a generalized Harmonic number, defined as

$$H_{x,p} = \sum_{m=1}^{x} m^{-p} = \zeta(p) - \sum_{m=x+1}^{\infty} m^{-p}$$

where $\zeta$ is the Riemann zeta function. Performing the Euler-Maclaurin expansion on the "tail" sum, we have

$$S_q(x) \approx \zeta\left(\frac{3}{2}\right) - q\zeta\left(\frac{5}{2}\right) - 2x^{-1/2} + \frac{3+4q}{6} x^{-3/2} + O(x^{-5/2}) \qquad (q < 1 < x) \qquad \text{(S9)}$$

For $q > 1$, we begin by performing the Euler-Maclaurin expansion directly on the original sum:

$$\begin{aligned} S_q(x) =& \sqrt{\frac{\pi}{q}} \left(\text{erf}(\sqrt{q}) - \text{erf}(\sqrt{q/x})\right) + \frac{e^{-q} + x^{-3/2}e^{-q/x}}{2} \\ &+ \frac{1}{12}\left(\frac{e^{-q/x}(-3x+2q)}{2x^{7/2}} - \frac{e^{-q}(-3+2q)}{2}\right) + ... \end{aligned} \tag{S10}$$

In the regime $q > x > 1$, we Taylor expand in small $x/q$ (for the $x$-dependent terms) and in small $1/q$ (for the terms independent of $x$). Our result is

$$\begin{aligned} S_q(x) =& e^{-q}\left(\frac{-q}{12} + \frac{5}{8} - \frac{1}{q} + ...\right) \\ &+ e^{-q/x}\left(\frac{\sqrt{x}}{q} + \frac{1}{2x^{3/2}} - \frac{x^{3/2}}{2q^2} + \frac{q}{12x^{7/2}} - \frac{1}{8x^{5/2}} + ...\right) \qquad (q > x > 1) \end{aligned} \tag{S11}$$

Since $x > 1$, the $e^{-q}$ terms are dwarfed by the $e^{-q/x}$ terms. For biologically reasonable parameters ($x < q < 100$), the first term is typically sufficient for a good fit.

In the regime $x > q > 1$ we start with the same Euler-Maclaurin expansion (Eq. S10), but rather than taking the Taylor series, we take the Bürmann expansion of $\text{erf}(\sqrt{q/x})$ for small $q/x$ in order to get faster convergence. The Bürmann series of the error function is

$$\text{erf}(z) = \frac{2\ \text{sgn}(z)}{\sqrt{\pi}}\left(\sqrt{1-e^{-z^2}}\right)\left(1 - \frac{1}{12}(1-e^{-z^2}) - \frac{7}{480}(1-e^{-z^2})^2 + O\left((1-e^{-z^2})^3\right)\right)$$

For the level of accuracy required for our work, it is sufficient to keep only the first term, yielding

$$\text{erf}(\sqrt{q/x}) \approx \sqrt{1-e^{-q/x}}$$

We then Taylor expand in small $x/q$ which converts this resulting exponential into a power series. As above, we neglect terms proportional to $e^{-q}$. The result is:

$$S_q(x) \approx \sqrt{\frac{\pi}{q}} - 2x^{-1/2} + \frac{3+4q}{6}x^{-3/2} + O(x^{-5/2}) \qquad (x > q > 1) \tag{S12}$$

In Supp Fig. 4, we plot $S_q(x)$ alongside these approximations (Eqs. S9, S11, S12).

### Section S4: A search using ITs

**The choice of $n$.** As discussed in the main text, each IT step takes the protein to a site a distance $m$ away with probability that scales as $m^{-3/2}e^{-q/m}$. The heavy tail of this probability distribution makes ITs superdiffusive. The typical time it takes to traverse a distance of order $L$ through ITs scales with $L$ as $L^{1/2}$ for large $L$. This implies that a search using ITs is limited not by the time it takes to reach the vicinity of the target site but by the time it takes to search locally near the target site. A search using ITs will often "leap" over the target site since large steps are common. Because of this, the search time with ITs is well described by considering ITs as taking the protein to a random (and uniformly chosen) site along the RNA. This same approximation has been made in other work on ITs as well [1]. To first order, therefore, the number of sites searched by the protein through ITs is given by the number of ITs taken (plus 1, for the site of initial non-specific association).

There are also two factors of 2 in the calculation of $n$ that we have neglected for simplicity. The first factor of 2 arises from that each time the protein is transiently bound at two

RNA segments, it has a 50% chance of dissociating from each segment before the other, and therefore only a 50% chance of transferring between the segments. The second factor of 2 arises from Cas13 having only one recognition domain (the crRNA), so if an IT brings the protein to the target sequence, there is a 50% chance of the protein not recognizing the target (since there is a 50% chance that the site at which the protein is bound to the target is the recognition domain). These factors of 2 affect the factor of $n$ in the search time (though they do not affect the dissociation rate), but do not have any qualitative effect on our results.

**Edge effects.** When the protein is at site $i$, it can transfer to another site $j$ or dissociate. The rate of transferring from site $i$ to site $j$ is given by $k_{ij}$, and the rate of dissociation is given by $k_{\mathrm{IT}}$. Finally, there is a site-independent rate $k_{\mathrm{on}}$ of binding at site $i$. The transfer rate matrix $k_{ij}$ is symmetric, and is defined as

$$k_{ij} = \alpha |i-j|^{-3/2} e^{-q/|i-j|}$$

The probability of the protein successfully transferring to any site $j$ through ITs, rather than dissociating, is given by

$$T_i = \frac{\sum_j k_{ij}}{\sum_m k_{i,m} + k_{\mathrm{IT}}}$$

Our goal in this section is to estimate $p_i$, the steady-state occupancy probability of the protein being at site $i$. At steady-state, the rate of entering state $i$ must equal the rate of leaving state $i$:

$$\sum_j k_{ji}\, p_j + k_{\mathrm{on}} = p_i \left( \sum_j k_{ij} + k_{\mathrm{IT}} \right)$$

or, using the symmetry of $k_{ji} = k_{ij}$,

$$p_i = \frac{\sum_j k_{ij} p_j + k_{\mathrm{on}}}{\sum_m k_{im} + k_{\mathrm{IT}}} \approx \sum_j \frac{k_{ij}}{\sum_m k_{im} + k_{\mathrm{IT}}} p_j$$

where we've assumed that $k_{\mathrm{on}}$ is negligibly small compared to the internal transfer dynamics.

This equation defines a recurrence relation for the probability vector $p$. By iterating this relation, we can estimate $p_i$. Starting from the uniform vector, the first iteration yields

$$p_i \propto \frac{\sum_j k_{ij}}{\sum_m k_{im} + k_{\mathrm{IT}}}$$

In other words, the steady-state occupancy probability of site $i$ is approximately proportional to $T_i$, the probability of performing a successful IT from site $i$ rather than dissociating. This leads to the edge effects shown in Fig. 3G.

When considering $n^{\mathrm{IT}}$, we neglect these edge effects, and assume $p_i$ is a constant. This enables us to write $n^{\mathrm{IT}} \approx k_R^{\mathrm{avg}}/k_{\mathrm{IT}}$ as discussed in the main text.

**A limit on $k_{\mathrm{IT}}$.** As noted in the main text, ITs are only useful if the RNA is able to re-equilibrate between each IT. The equilibration time can be approximated as the time it takes for an RNA segment to travel a distance equal to the size of the molecule. The end-to-end monomer motion within the coil scales as [12]

$$\sqrt{\langle r^2 \rangle} = \sqrt{2} \left( \frac{k_B T}{\eta} t \right)^{1/3}$$

The equilibration time therefore scales as the cube of the radius of gyration of the RNA. As discussed above, for RNA, $r \approx 0.5\ L^{1/3}$ nm, meaning that the equilibration time is approximately given by

$$t_{\mathrm{equil}} \approx 2^{-9/2}\ \mathrm{nm}^3 \frac{\eta}{k_B T} L \approx 10^{-10} L\ \mathrm{s}$$

In other words, for an RNA of length $L = 10^3$, the equilibration time is approximately equal to $10^{-7}$ s. This sets an approximate upper limit for the rate of ITs: ITs that occur faster than this would not allow the RNA to fully re-equilibrate in between each IT, leading to increased correlations between successive sites visited. This regime is not considered in this work. We find that for the parameter regimes considered, the optimal rate of ITs is slower than this limit, so we can assume the RNA re-equilibrates between each IT (Fig. 5).

### Section S5: The effects of Cas13 search time on phage infection

In Fig. 1B we show the results of a model for the effect of Cas13 on phage infection, including the role played by the search time. In this section, we describe the model.

We describe the phage infection process and Cas13's role through a system of ordinary differential equations (ODEs). We consider a bacterial cell that contains crRNA/Cas13 complexes. The cell may contain many such complexes. At time $t = 0$ a DNA phage infects the cell, and begins transcribing mRNAs. We consider the case in which one of the mRNAs transcribed by the phage contains a protospacer labeling it as a target for one of the crRNA/Cas13 complexes in the bacterial cell. We denote by $N$ the number of Cas13 enzymes complexed to crRNAs that target this mRNA. For simplicity, we will simply refer to these as Cas13, since no other Cas13 enzymes in the cell are relevant. We neglect Cas13/crRNA dissociation, and assume that the bacterial cell has only one crRNA species able to target the phage.

We assume that the relevant mRNA is transcribed at a constant rate $\alpha$, and is degraded at a basal rate $\mu_0$. We also assume the mRNA is translated into a protein $H$ which leads to lysis when it accumulates past a certain threshold $H^*$. The rate of translation is denoted by $k_h$.

Cas13 encounters the mRNA with a bimolecular association rate $k_2$. However, since the number $N$ of searching Cas13 molecules is small, we consider Cas13 activation as a stochastic process, which we describe later. The number of active Cas13 enzymes is denoted by $A$. Cas13 activation has two effects within this model: first, Cas13 cleaves the mRNAs, increasing the mRNA degradation rate by an amount $\mu_A$. Second, Cas13 cleaves other cellular RNAs including rRNAs, leading to a decrease in the overall transcription rate $\alpha$. The rate of this decrease is denoted by $\delta$. The equations governing the system behavior are given below:

$$\frac{dM}{dt} = \alpha - (\mu_0 + \mu_A A)M \tag{S13}$$

$$\frac{dH}{dt} = k_h M \tag{S14}$$

$$\frac{d\alpha}{dt} = -\delta A \alpha \tag{S15}$$

We now describe the stochastic process by which we model Cas13 activation. To efficiently simulate the low-copy-number activation of Cas13 alongside the high-copy-number accumulation of mRNA and lytic proteins, we employed a Piecewise Deterministic Markov Process (PDMP) framework. In particular, while the ODEs above are integrated using the Euler method (with $\Delta t = 0.01$ min), Cas13 activation is modeled stochastically using a time-dependent hazard integration method. The instantaneous activation propensity is given by

$$\frac{dQ}{dt} = k_2 M(N - A) \tag{S16}$$

At the start of the simulation (or immediately following an activation event), $Q$ is set to zero and a random threshold $Q^*$ is sampled from a standard exponential distribution. The integrated hazard $Q(t)$ is updated at each time-step using the Euler method, as for $M$, $H$, and $\alpha$. An activation event is triggered when $Q \geq Q^*$, at which point $Q$ is reset to zero, a new $Q^*$ is drawn, and $A$ is incremented by one. This hybrid approach captures the stochasticity of rare molecular events without the prohibitive computational cost of a full Gillespie simulation.

We now describe the stochastic process by which we model Cas13 activation. To efficiently simulate the low-copy-number activation of Cas13 alongside the high-copy-number accumulation of mRNA and lytic proteins, we employed a Piecewise Deterministic Markov Process (PDMP) framework [13]. In particular, while the ODEs above are integrated using the Euler method (with $\Delta t = 0.01$ min), Cas13 activation is modeled stochastically using a time-dependent hazard integration method. The instantaneous activation propensity is given by

$$\frac{dQ}{dt} = k_2 M(N - A) \tag{S17}$$

At the start of the simulation (or immediately following an activation event), $Q$ is set to zero and a random threshold $Q^*$ is sampled from a standard exponential distribution. The integrated hazard $Q(t)$ is updated at each time-step using the Euler method, as for $M$, $H$, and $\alpha$. An activation event is triggered when $Q \geq Q^*$, at which point $Q$ is reset to zero, a new $Q^*$ is drawn, and $A$ is incremented by one. This hybrid approach captures the stochasticity of rare molecular events without the prohibitive computational cost of a full Gillespie simulation.

We now describe the stochastic process by which we model Cas13 activation. To efficiently simulate the low-copy-number activation of Cas13 alongside the high-copy-number accumulation of mRNA and lytic proteins, we employed a Piecewise Deterministic Markov Process (PDMP) framework. In particular, while the ODEs above are integrated using the Euler method (with $\Delta t = 0.01$ min), Cas13 activation is modeled stochastically using a time-dependent hazard integration method. The instantaneous activation propensity is given by

$$\frac{dQ}{dt} = k_2 M(N - A) \tag{S18}$$

At the start of the simulation (or immediately following an activation event), $Q$ is set to zero and a random threshold $Q^*$ is sampled from a standard exponential distribution. The integrated hazard $Q(t)$ is updated at each time-step using the Euler method, as for $M$, $H$, and $\alpha$. An activation event is triggered when $Q \geq Q^*$, at which point $Q$ is reset to zero, a new $Q^*$ is drawn, and $A$ is incremented by one. This hybrid approach captures the stochasticity of rare molecular events without the prohibitive computational cost of a full Gillespie simulation.

The stochasticity in $Q^*$ leads to a stochasticity in the results shown in the inset to Fig. 1B. For the same parameter set, different iterations of the simulation lead to different results. Essentially, the larger $k_2$ is (i.e. the shorter Cas13's search time), the more consistently the bacterial cell is able to thwart the phage infection (keeping $H < H^*$). $k_2$ is related to the search time by $(t_{\text{search}})^{-1} = k_2(1\ \mu\text{m}^{-3})$; i.e. the search time is the rate of a single Cas13 enzyme finding a single mRNA within the cellular volume.

### Parameters used in Fig. 1B:

- $N = 2$
- $\alpha(t = 0) = 2.4$ mRNAs/cell/min (estimated with a transcription rate of 40 nt/s (BNID[6] 106941) and an mRNA length of $10^3$ nts).
- $H^* = 10^3$ (chosen such that the lysis time in the absence of Cas13 is approximately 20 min).
- $\mu_0 = \ln(2)/5$ min (BNID: 111927)

- $\mu_A = 0.5/\text{min} = 3.6\ \mu_0$
- $k_h = 5/\text{min}$ (BNID: 112001)
- $\delta = 0.01/\text{min}$

Parameters used in other figures are given in Supp. Table 2. For DNA, ITs are modeled as for RNA, with $q = 0$ and $\alpha$ smaller by a factor of $10^5$ as discussed in the main text.

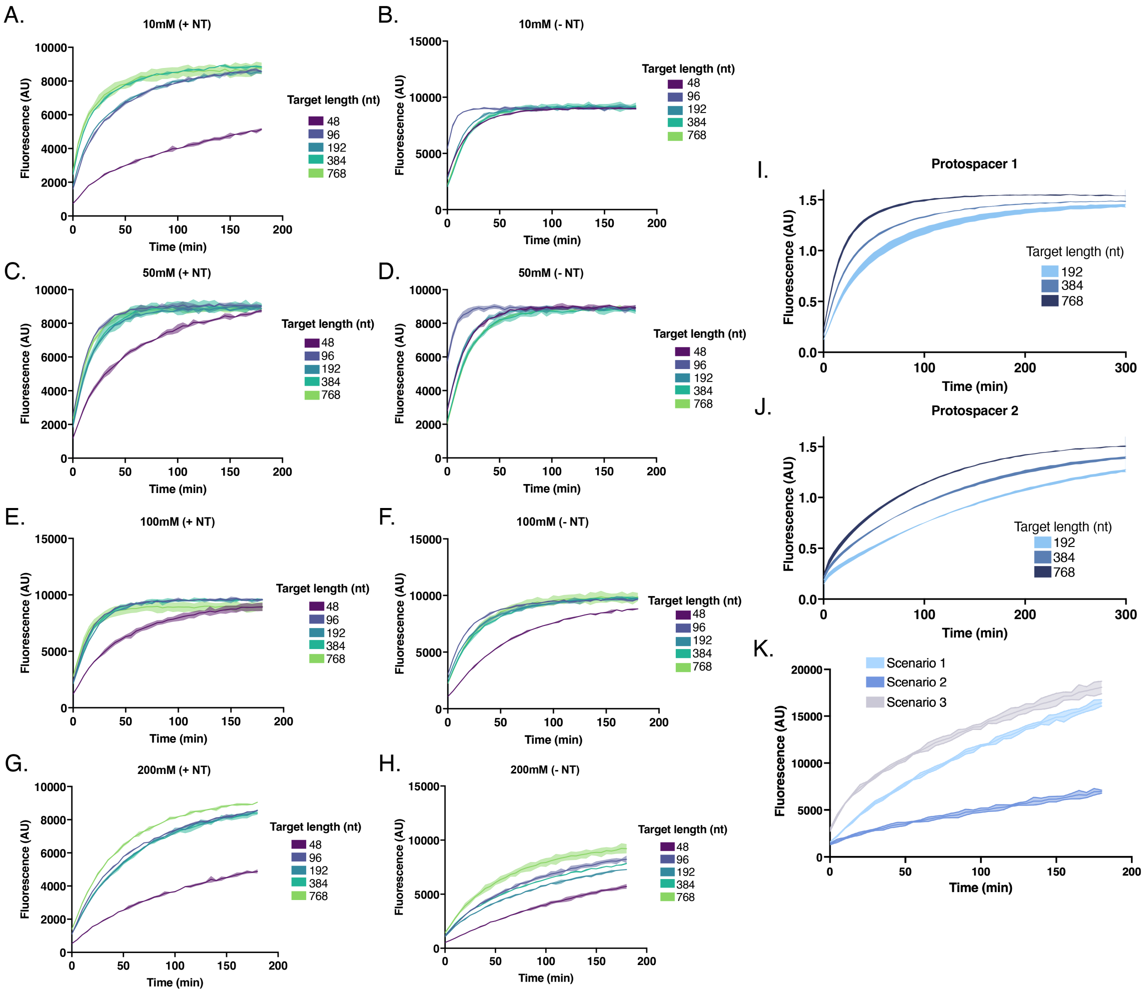


**Supplementary Figure 1. Raw Cas13 detection curves for ionic concentration, target length, and intersegmental transfer assays. A.** Detection curves for Cas13 assay targeting different length RNAs in a buffer containing 10 mM NaCl, in the presence of non-target RNA. **B.** Curves as in A, but in the absence of non-target RNA. **C-H.** As in A-B, but in buffers containing increasing concentrations of NaCl. **I-J.** Detection curves for Cas13 assays targeting RNAs of three different lengths, for two different protospacer sequences. **K.** Detection curves for the intersegmental transfer assay described in Fig. 4A. Shading shows error of n = 2 technical replicates.

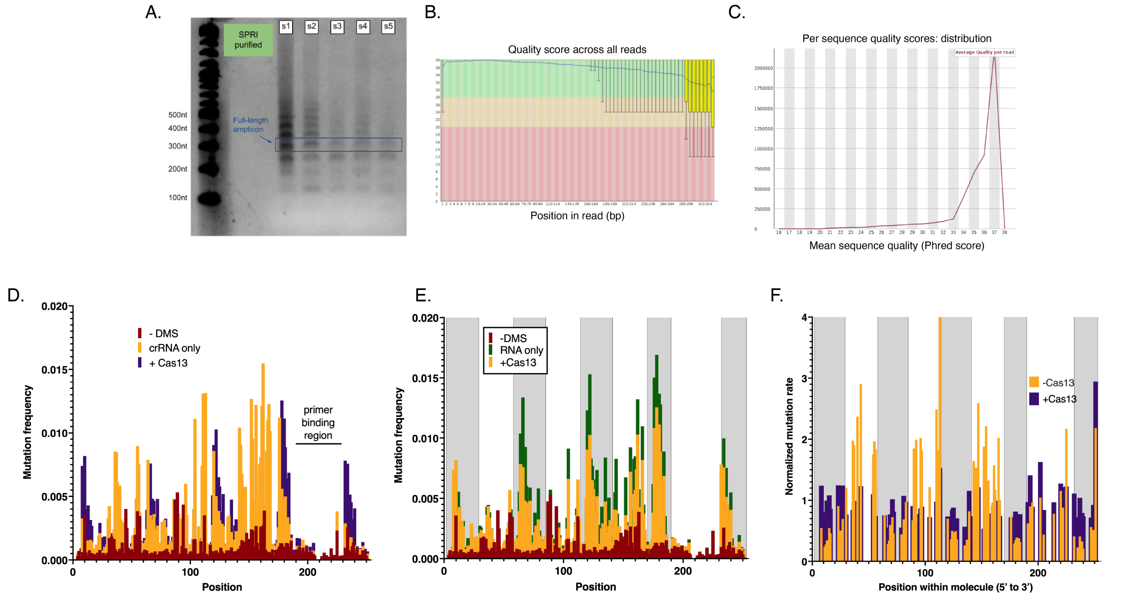


**Supplementary Figure 2. Sequencing quality control and raw data from DMS footprinting assay. A.** Representative agarose gel showing the amplicon to be sequenced following library preparation, along with expected truncated or concatemer products owing to the RNA circularization step. **B.** Per-base sequence quality scores across the length of the amplicon. Generated using MultiQC. **C.** Read quality histogram. Generated using MultiQC. **D.** Raw mutation frequencies for the untreated control, crRNA-only, and full RNP conditions are shown across the complete length of the RNA molecule. **E.** As in D, but comparing untreated, RNA only, and full RNP conditions. **F.** Comparing mutation frequency of A bases for the -Cas13 and +Cas13 conditions only. If E and F, protospacer sequences are shaded gray.

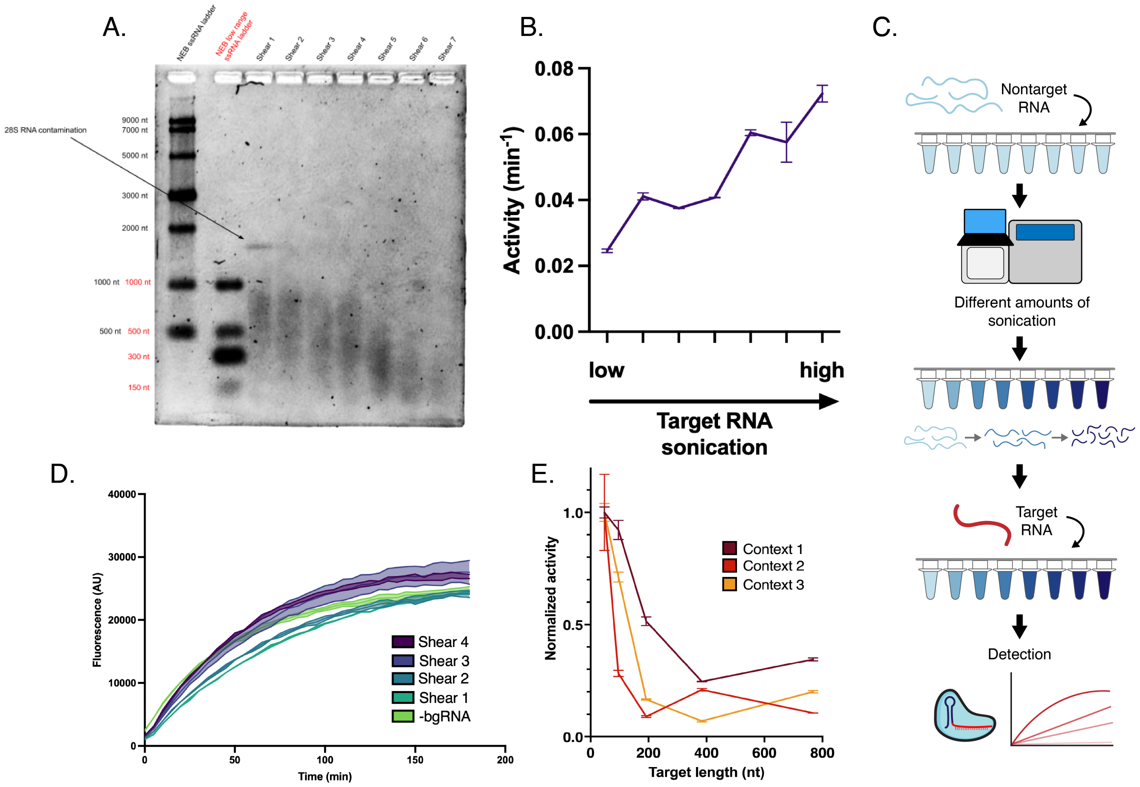


**Supplementary Figure 3. Experiments involving target and non-target RNA sonication.** **A.** Agarose gel electrophoresis of sonicated RNA samples confirming average RNA length. **B.** Activity scores from Cas13 detection assays when targeting differently sonicated RNAs from A. Error bars represent the standard deviation of two technical replicates. **C.** Schematic describing the workflow used to assess the impact of different non-target RNA lengths using sonication. **D.** Raw curves from Cas13 detection assay with differently sheared non-target RNAs. Shading shows error of n = 2 technical replicates. **E.** Activity scores from Cas13 detection assays targeting RNAs of differing lengths with the same protospacer sequence but different sequence contexts surrounding the protospacer. Error bars represent the standard deviation of two technical replicates.

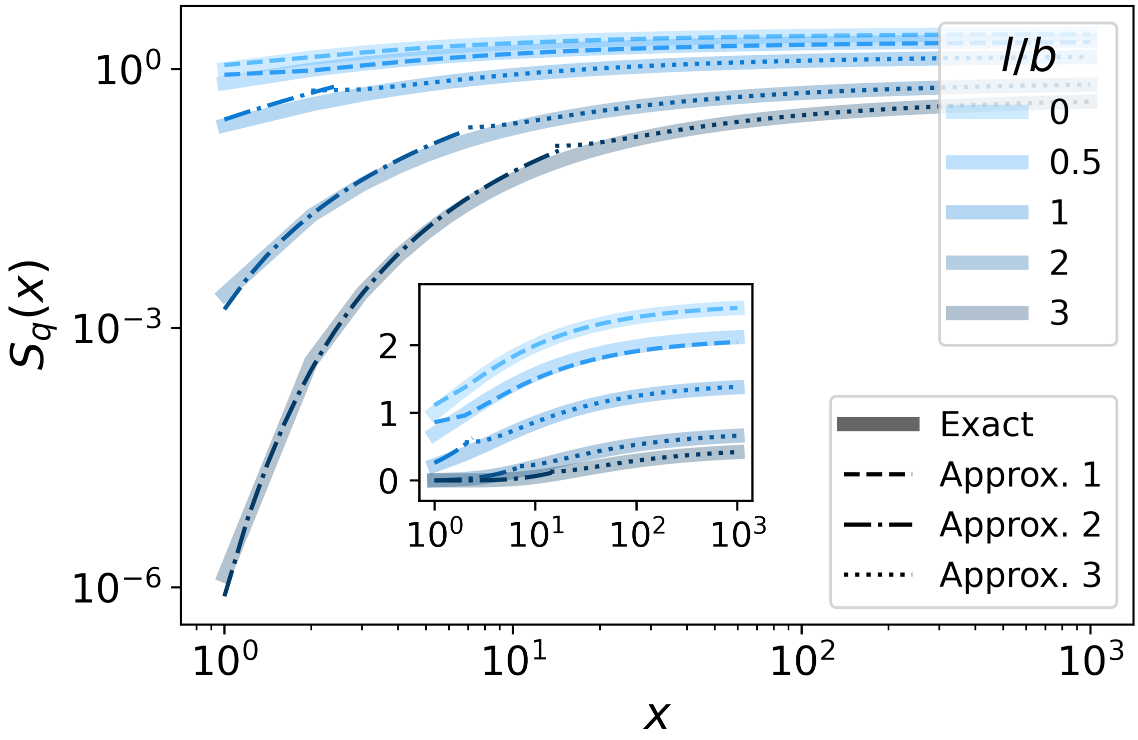


**Supplementary Figure 4. Scaling of $S_q(x)$ as a function of relative protein position $x$.** $S_q(x)$ is plotted for various values of $q = 3l^2/2b^2$ (different colors). Lines show approximate forms of the sum derived in Supp. Section S3. Dashed lines (approx. 1) plots $\zeta(3/2)-q\zeta(5/2)-2x^{-1/2}$ (Eq. S9; valid for $q < 1$). Dash-dotted lines (approx. 2) show $e^{-q/x}(x^{1/2}q^{-1}+x^{-3/2}/2)$ (Eq. S11; valid for $q > x > 1$). Dotted lines (approx. 3) show $(\pi/q)^{1/2} - 2x^{-1/2} + (3 + 4q)x^{-3/2}/6$ (Eq. S12; valid for $x > q > 1$).

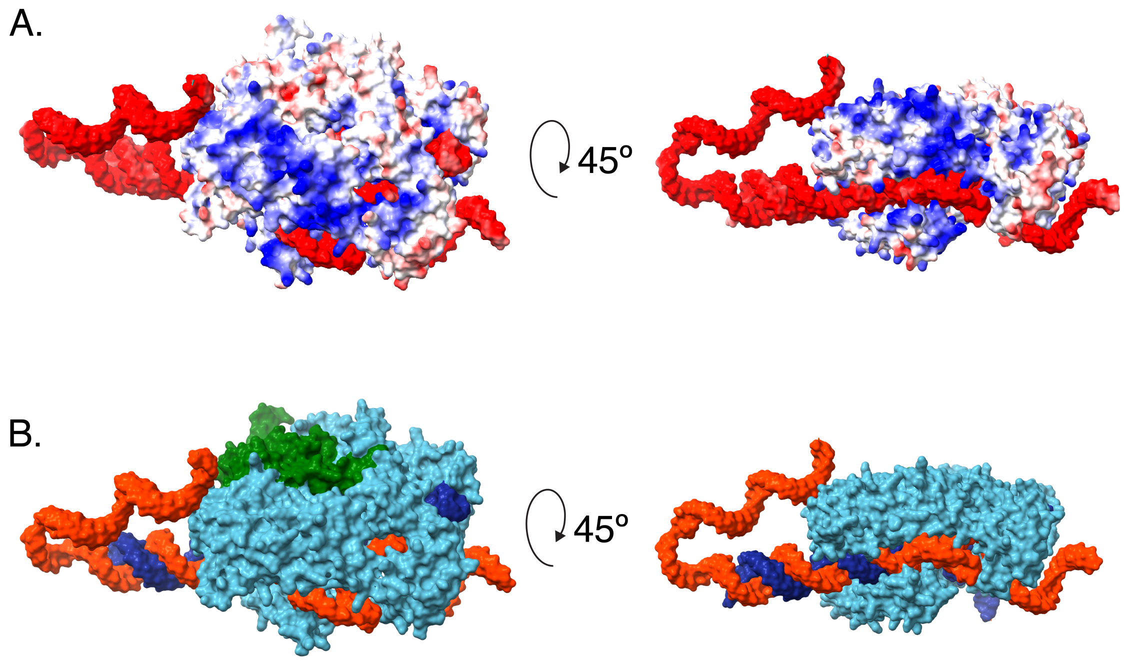


**Supplementary Figure 5. AlphaFold 3 renderings of target-bound LwaCas13a. A.** Two views of the target-bound LwaCas13a complex, with the atoms colored by electrostatic potential (red = negatively charged, blue = positively charged). **B.** The same views as in A, colored to show the different molecules and domains. Orange: target RNA; dark blue: crRNA; light blue: Cas13; green: HEPN domain. In A and B, AlphaFold 3 renderings were generated using UCSF ChimeraX.

[2] Marian Smoluchowski. Versuch einer mathematischen Theorie der Koagulationskinetik kolloider Lösungen. *Zeitschrift für Physikalische Chemie*, 92, 1917.
